# Transcytosis-mediated anterograde transport of TrkA receptors controls formation of presynaptic sites in sympathetic neurons

**DOI:** 10.1101/2025.09.18.677078

**Authors:** Guillermo Moya-Alvarado, Sebastian M Markert, Sumana Raychaudhuri, Matthew Tachoute, Shigeki Watanabe, Rejji Kuruvilla

## Abstract

In neurons, many membrane proteins, synthesized in cell bodies, must be efficiently delivered to axons to influence neuronal connectivity. Transcytosis is an atypical transport mode, where membrane proteins internalized from soma surfaces are anterogradely transported to axons. Here, we define the trafficking dynamics and the transport vesicles involved in transcytosis of TrkA neurotrophin receptors and demonstrate that transcytosis controls formation of presynaptic sites in sympathetic neurons. Live imaging and electron microscopy in compartmentalized cultures revealed that soma surface-derived TrkA receptors undergo dynamic movements in axons and are housed in endosomes and multi-vesicular bodies. Soma-surface labeled TrkA appear in nerve terminals, demonstrating transcytosis occurs *in vivo*. Notably, transcytosed TrkA receptors are enriched at presynaptic varicosities. Disruption of transcytosis impairs the number and morphology of presynaptic sites and decreases synaptic transmission. These findings provide mechanistic insight into an atypical mode of receptor trafficking and highlights its physiological relevance in sympathetic neuron connectivity.

## Introduction

The immense length of axons imposes unique challenges on neurons in controlling cellular functions in distal axon compartments. Axon terminals can be meters away from cell bodies where many axonal membrane proteins with critical functions in regulating axon guidance and growth, neuronal survival, presynaptic organization, and synaptic transmission, are made. These proteins need specialized mechanisms to be transported to their final destinations either via direct trafficking through the secretory pathway after sorting at the trans-Golgi network or transcytosis ^1,2^. Transcytosis is an atypical endocytosis-based mechanism, where newly-synthesized proteins are first inserted on cell body surfaces, internalized, and anterogradely transported to axons. In contrast to the considerable progress made in understanding the direct secretory pathway, there is limited knowledge about transcytosis, specifically, the underlying transport kinetics and organelles involved, whether it occurs in *vivo*, and its contributions to neuronal connectivity and function.

The family of tropomyosin-related kinases (Trk) receptors provides a prominent example of membrane proteins that undergo long-distance axonal trafficking to control neuronal survival, axon growth, and synaptic transmission ^3, 4^. In sympathetic and sensory neurons, axonal TrkA receptors are internalized after binding the ligand, Nerve Growth Factor (NGF), secreted from peripheral tissues ^5, 6^. Axon-derived TrkA receptors are then retrogradely transported long-distance to neuronal cell bodies to exert transcriptional control of developmental programs. However, neuronal responsiveness to target-derived NGF also requires the precise axonal targeting of new TrkA receptors, synthesized in cell bodies. Previously, we discovered that newly synthesized TrkA receptors are delivered to axons of sympathetic neurons by transcytosis ^7, 8^. Remarkably, in contrast to the constitutive secretory pathway, anterograde TrkA transcytosis is regulated by the ligand, NGF, acting on distal axons ^7, 8^. Since NGF is a target-derived trophic factor for sympathetic neurons and is secreted in limiting amounts during development, regulated transcytosis suggests a positive feedback mechanism that serves to dynamically scale up receptor availability in axons at times of need. Together, these previous findings provide an entry point to investigate a poorly characterized ligand-promoted mode of axonal targeting of membrane proteins. Several outstanding questions remain about the transport kinetics of TrkA transcytosis, the identity of the organelles responsible for receptor transcytosis, and its functional relevance in the sympathetic nervous system.

Here, we define the dynamic behavior of transcytosing TrkA receptors, identify the organelles involved, and demonstrate that TrkA transcytosis is necessary for the formation and function of presynaptic varicosities in sympathetic axons. Using live imaging in compartmentalized neuron cultures, we observed soma surface-derived TrkA exhibiting distinct behaviors upon transcytosis to distal axons, with anterograde movements, pausing, and intriguingly, even retrograde movements. Electron microscopy revealed that transcytosing TrkA receptors are localized primarily in endosomes and multivesicular bodies in proximal axon compartments but transition to a predominantly endosomal localization in distal axon compartments. Using a paradigm to label soma surface TrkA receptors in sympathetic ganglia, we found that soma surface-derived TrkA receptors appear in nerve terminals *in vivo*, establishing receptor transcytosis as a physiological mode of axon targeting. Notably, transcytosed TrkA receptors are localized to presynaptic axonal varicosities. Disruption of TrkA transcytosis impaired the number and morphology of presynaptic sites *in vitro* and *in vivo*, as well as decreased synaptic transmission between sympathetic neurons and cardiomyocytes in co-cultures. These findings provide mechanistic insight into an atypical mode of anterograde receptor transport and highlights the physiological relevance of TrkA transcytosis in sympathetic neuron connectivity and function.

## Results

### Live imaging of soma surface-derived TrkA receptors reveals distinct behaviors in proximal *versus* distal axons

To obtain a dynamic view of the behavior and transport kinetics of TrkA receptors undergoing transcytosis, we performed live imaging of cultured sympathetic neurons obtained from a *Ntrk1^Flag^* knock-in mouse line, which expresses Flag epitope-tagged TrkA protein from the endogenous TrkA locus ^9^. Neurons were grown in compartmentalized microfluidic chambers, and soma surface Flag-TrkA receptors were live-labeled exclusively in cell body compartments with anti-Flag antibodies conjugated with the Cy3 fluorophore (**Fig. 1A**). Application of the ligand, NGF, exclusively to distal axon compartments allows for real-time visualization of anterograde transcytosis of soma-surface TrkA receptors to axons. In the absence of NGF, the majority of Flag-TrkA puncta were stationary in proximal axon compartments (**Fig 1B, D**). After 30 min of stimulation of distal axons with NGF, we observed a pronounced increase in the movement of soma surface-derived Flag-TrkA receptors in the anterograde direction in proximal axons (**Fig. 1C, D**). Kinetic analyses revealed that 25.1 ± 2.97% of Flag-TrkA^+^ particles were anterogradely transported in proximal axons, with an average instant speed of 1.02 ± 0.06 µm/s, while the majority of particles (72.6 ± 2.94%) were stationary (**Fig. 1D, E and Supplementary movie 1**). After 30 min of NGF treatment, there were little to no soma surface-derived TrkA puncta appearing in distal axon compartments. However, after 180 min of NGF stimulation of distal axons, Flag-TrkA receptors, initially labeled on soma surfaces, were observed in distal axons, consistent with anterograde transcytosis (**Fig. 1F**). Upon arrival in distal axons, transcytosed TrkA receptors exhibited distinctive behaviors (**Fig. 1F**), with 16.3 ± 2.5% of particles moving anterogradely, 53.4 ± 3.9% remaining stationary, and intriguingly, a considerable population (25.9 ± 4.1%) moving in the retrograde direction (**Fig. 1F, G and Supplementary movie 2**). The instant speed of anterogradely moving particles (0.38 ± 0.05 µm/s) was significantly slower than that of the retrogradely transported particles (1.18 ± 0.08 µm/s) (**Fig. 1H**), which could reflect differences in intrinsic properties of molecular motors used for anterograde (kinesins) versus retrograde transport (dynein), as well as differences in interactions of motors with adaptors, microtubules, or axonal organelles ^10^.

**Figure 1.**
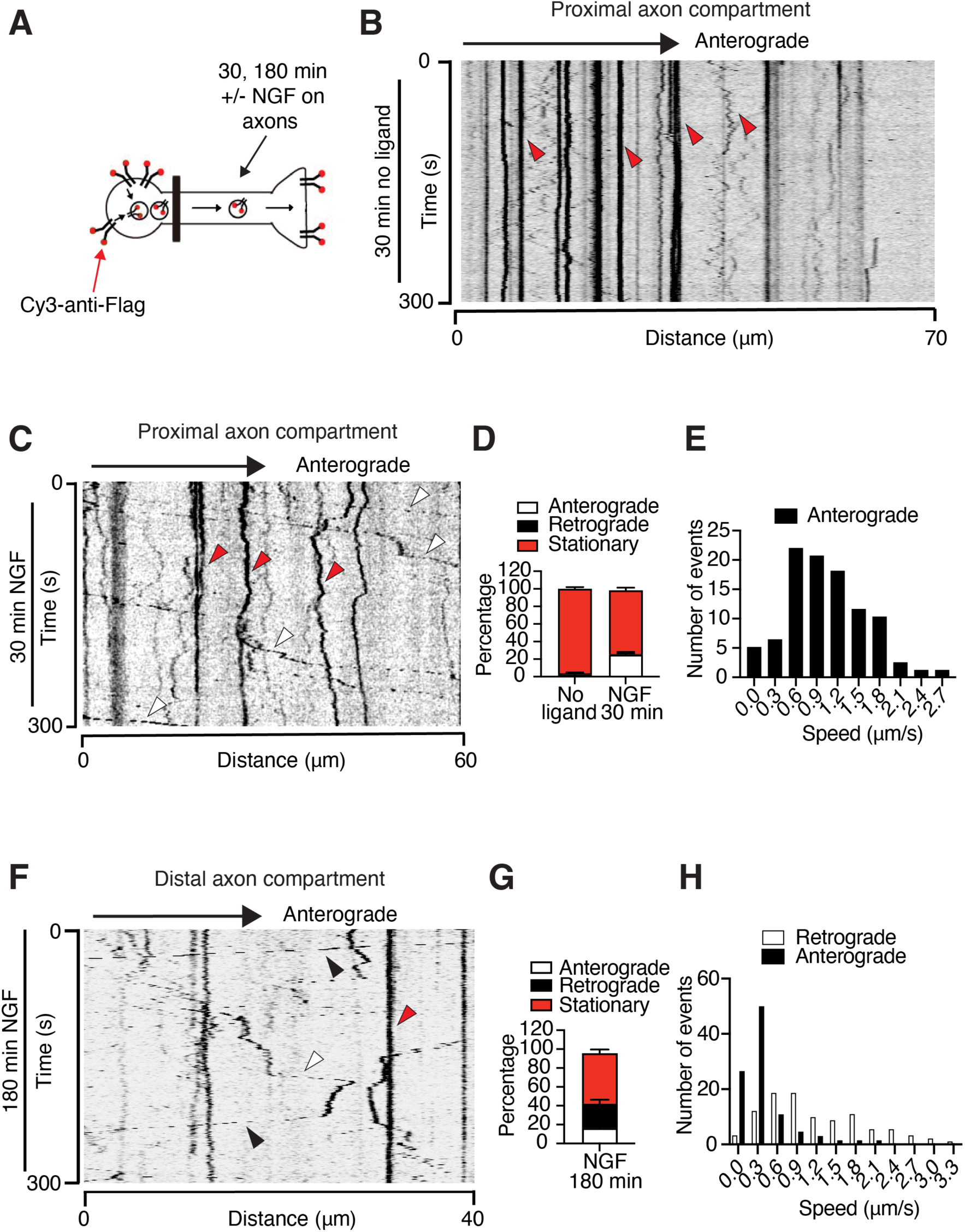
Live imaging of TrkA transcytosis. (**A**) Schematic of antibody feeding assay to monitor Flag-TrkA transcytosis using live imaging in Ntrk1^Flag^ mouse sympathetic neurons grown in microfluidic cultures. Soma surface Flag-TrkA receptors were live-labeled by feeding Cy3-anti-Flag antibodies to soma + proximal axon compartments. Distal axons were either left untreated or stimulated with NGF (100 ng/mL) for 30 or 180 min. (**B**) Kymograph showing behavior of Flag-TrkA particles in proximal axons in the absence of ligand. (**C**) Kymograph showing dynamic behavior of Flag-TrkA particles in proximal axons after NGF stimulation of distal axons (30 min). (**D-E**) Quantification of directionality (**D**) and instant speed (**E**) of Flag-TrkA particles in proximal axons. (**F**) Kymograph showing behavior of Flag-TrkA particles in distal axons after NGF stimulation of distal axons (180 min). (**G-H**) Quantification of directionality (**G**) and instant speed (**H**) of Flag-TrkA in distal axons. Data are means ± SEM from 4 independent experiments (total of n=21 kymographs analyzed).

To compare the transport dynamics of TrkA transcytosis with retrograde trafficking of axon-derived TrkA receptors, we incubated distal axon compartments of *Ntrk1^Flag^* sympathetic neuron cultures with Cy3-conjugated anti-Flag antibodies, followed by NGF stimulation of distal axons for 30 min (**Supplementary Fig. 1A**). Live-imaging of axon-derived TrkA receptors showed that the majority (74.9 ± 9.6%) undergo highly processive movement almost exclusively in the retrograde direction (**Supplementary Fig. 1B**), with a net speed of 1.59 ± 0.10 µm/s (**Supplementary Fig. 1C**), consistent with previous studies ^11, 12^.

These findings indicate that the behavior and dynamics of movement of soma surface-derived TrkA receptors undergoing transcytosis differ markedly within proximal *versus* distal axon compartments, as well, as from the retrograde transport of axon-derived TrkA receptors.

### Ultrastructural analysis of TrkA transcytosis

We next used electron microscopy (EM) to define the ultrastructural features of organelles carrying soma surface-derived TrkA receptors undergoing anterograde transcytosis. For this purpose, cell body compartments of *Ntrk1^Flag^* sympathetic neurons, grown in compartmentalized microfluidic cultures, were incubated with anti-Flag antibodies pre-conjugated to anti-rabbit IgG (H+L)-10 nm gold particles for 30 min (**Fig. 2A**). Distal axon compartments were then stimulated with NGF for 180 min (**Fig. 2A**) followed by EM imaging. The antibody labeling strategy revealed gold-labeled Flag-TrkA receptors, originating from soma surfaces, distributed in proximal as well as distal axon compartments. Of note, little to no electron-dense structures were observed in *Ntrk1^Flag^*sympathetic neuron cultures incubated with anti-rabbit IgG-10 nm gold particles or with addition of anti-Flag antibody-conjugated gold particles to wild-type sympathetic neuron cultures, pointing to the specificity of labeling (**Supplementary Fig. 2A, B)**.

**Figure 2.**
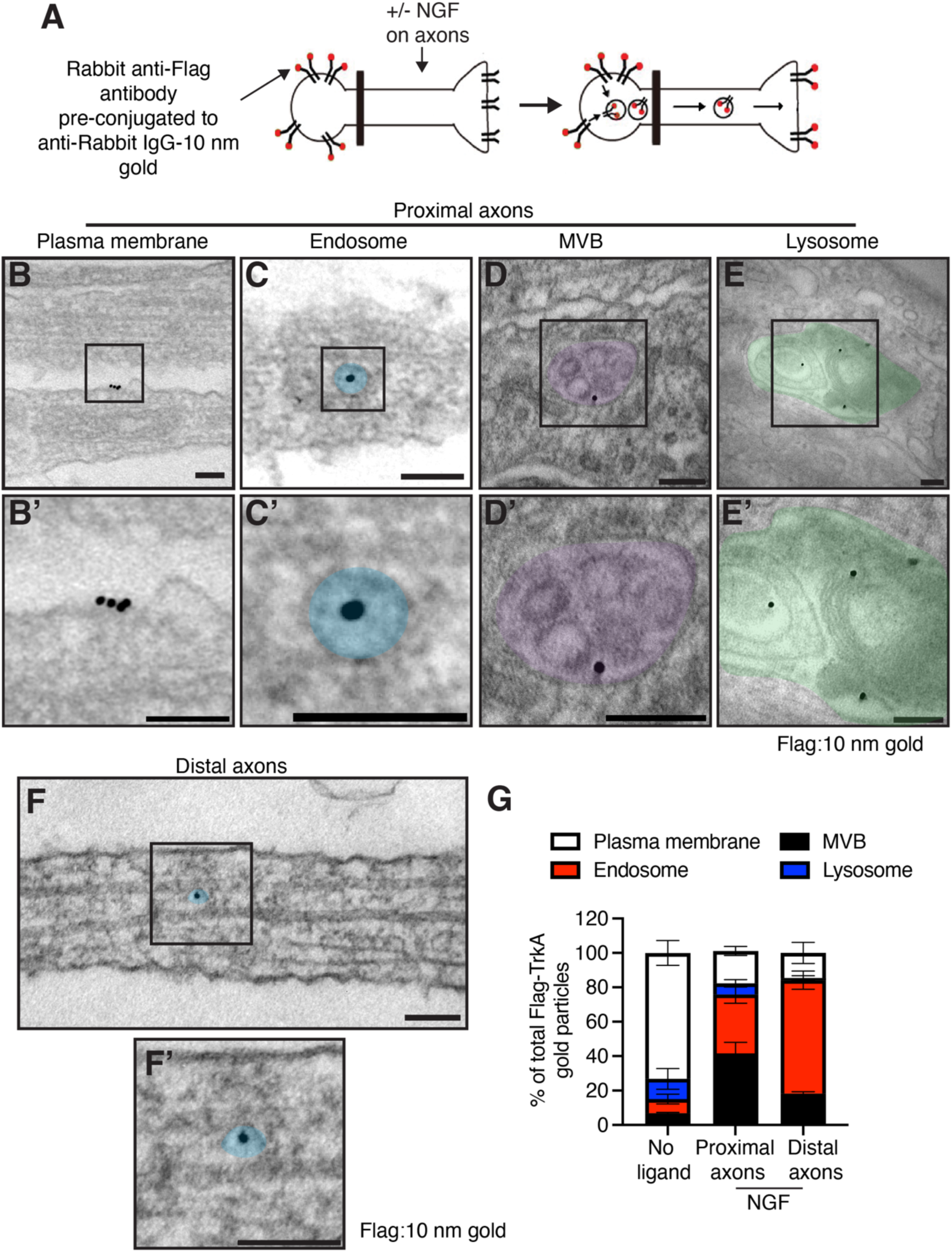
Ultrastructural analysis of TrkA transcytosis. (**A**) Schematic of antibody feeding assay to monitor Flag-TrkA transcytosis using electron microscopy in Ntrk1^Flag^ mouse sympathetic neurons grown in microfluidic cultures. Soma surface Flag-TrkA receptors were live-labeled with rabbit anti-Flag antibody pre-conjugated to anti-rabbit IgG 10 nm gold particles added specifically to soma + proximal axon compartments. Distal axons were either left untreated or stimulated with NGF (100 ng/mL) for 180 min. (**B-E’**) Representative images of Flag-TrkA gold particles at the plasma membrane (**B, B’**); within an endosome (**C, C’**), within a multivesicular body (MVB (**D, D’**)), or in a lysosome (**E, E’**). Higher magnification images of insets in top panels are shown in panels **B’-E’.** Scale: 100 nm. (**F, F’**) Representative image of Flag-TrkA gold particles in an endosome in the distal axon. Higher magnification image of inset in **F** is shown in **F’**. Scale: 100 nm. (**G**) Quantification of the percentage of Flag-TrkA gold-labeled particles located at the plasma membrane or within internal compartments in the absence of NGF as well as proximal and distal axons in NGF-stimulated neurons. Data are represented as means ± SEM from n=3 independent experiments.

In proximal axon compartments, Flag-TrkA gold particles were observed at the plasma membrane (**Fig. 2B, B’**), and in internal compartments with ultrastructural features, indicative of endosomes (single-membrane vesicular structures) (**Fig. 2C, C’**), multivesicular bodies (MVBs) (**Fig. 2D, D’**) and lysosomes (**Fig. 2E, E’**). Quantification of the distribution of Flag-TrkA gold particles in proximal axon compartments revealed that axonal NGF stimulation significantly shifted the pattern of receptor localization from being at the plasma membrane (73.2 ± 7.2% in the absence of NGF) to internal compartments with 34.2 ± 5.1% of receptors found in endosomes, 41.6 ± 6.4% in multivesicular bodies, and to a lesser extent in lysosomes (6.5 ± 2.1%) or still remaining at the plasma membrane (18.8 ± 2.6%) (**Fig. 2G**). Notably, in distal axon compartments, gold-labeled Flag-TrkA particles were predominantly localized in endosomes (66 ± 5.3 %) (**Fig. 2F, F’, G**), with a smaller population in multivesicular bodies (18.3 ± 1.1%), 14.5 ± 6.1% at the plasma membrane, and 1.3 ± 1.3% in lysosomes (**Fig. 2G**).

These results suggest that TrkA receptors undergoing transcytosis are primarily localized in endosomes and multivesicular bodies in proximal axon compartments but shift to an endosomal localization in distal axons. These findings imply a sorting of organelles carrying soma-surface receptors as they transit from proximal to distal axon compartments.

### TrkA receptors undergo transcytosis *in vivo*

To date, almost all information about axonal delivery of proteins has been obtained from culture studies. Little is known about physiological modes of axon delivery of membrane proteins. To assess TrkA transcytosis *in vivo*, we established a paradigm to label soma surface TrkA receptors residing in the superior cervical ganglia and to monitor the appearance of labeled receptors in nerve terminals innervating target tissues, including the salivary glands and iris, in living animals. Cell surface TrkA receptors in sympathetic ganglia were labeled by two independent approaches (**Fig. 3A**); (1) a biochemical approach by locally injecting membrane-impermeable biotin (sulfo-NHS-SS-biotin) into the superior cervical ganglion in rat pups (postnatal days 1-2), or (2) an immunohistochemical approach to visualize TrkA transcytosis by locally injecting anti-Flag antibodies into the superior cervical ganglion in *Ntrk1^Flag^* knock-in mice (postnatal days 1-2). Since the superior cervical ganglia are bilateral in the sympathetic chain, injections were done in one of each paired ganglia per animal, with the contralateral ganglion and target tissues (non-injected side) serving as internal controls to assess any systemic leakage of injected label. Of note, we used rat pups for surface biotinylation assays, since this approach yields more material for biochemical analyses compared to mice. After 8 hr, sympathetic ganglia and target tissues (salivary glands, iris) were dissected and subjected to either streptavidin pull-down/immunoblotting to detect biotin-labeled TrkA receptors or immunohistochemical analyses to visualize Flag-labeled receptors.

**Figure 3.**
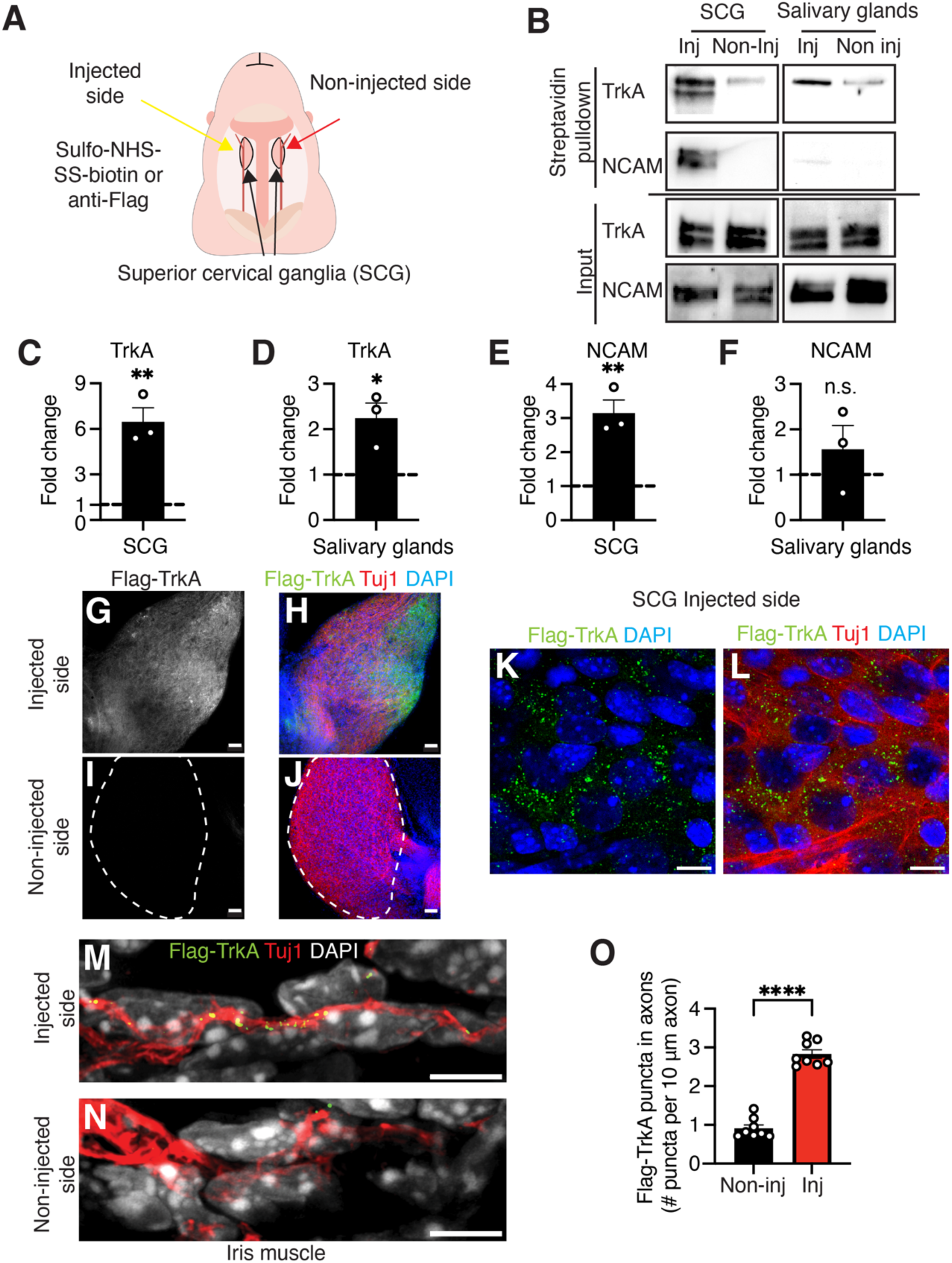
TrkA receptors are transcytosed from cell bodies to nerve terminals *in vivo*. (**A**) Schematic of *in vivo* injections of sulfo-NHS-SS-biotin or rabbit anti-Flag antibody to the superior cervical ganglia (SCG) in rats and Ntrk1^Flag^ mice, respectively, at P2-P3. Injections were done in one of each paired ganglia per animal with the contralateral ganglion and target tissues (non-injected side) serving as internal controls to assess any systemic leakage of injected label. (**B**) Immunoblotting for biotinylated TrkA and NCAM after streptavidin pull-downs in SCG and salivary gland lysates from injected and non-injected sides, 8 hr after sulfo-NHS-SS-biotin injection (top two panels). Bottom two panels show TrkA and NCAM immunoblotting in input lysates. (**C-F**) Increase in TrkA and NCAM levels in SCG and salivary glands at the injected side, relative to the non-injected side. Quantification is represented as fold-change relative to non-injected side (dashed line). Data are means ± SEM from n=3 animals; p*<0.05; **p<0.01, n.s: not significant, t-test (**G-J**) Flag and Tuj1 immunofluorescence in the SCG, 8 hr after injection of anti-FLAG antibodies in SCG. Injected side is shown in (**G,H**) and non-injected side in (**I,J**). Tuj1 (red), Flag-TrkA (gray (**G**), in green (**H**)), DAPI (Blue). Scale bar 50 μm. (**K-L**) Higher magnification image of injected SCG shows internalized Flag-TrkA puncta (green) in sympathetic neurons (Tuj1, red). DAPI is shown in blue. Scale bar 10 μm. (**M-N**) Increased Flag-TrkA puncta in axons innervating the iris at the injected side (**M**), compared to non-injected side (**N**) in Ntrk1^Flag^ mice. Tuj1 (red), Flag-TrkA (green), DAPI (gray). Scale bar 5 μm. (**O**) Quantification of Flag-TrkA puncta in axons innervating the irises from injected and non-injected sides, 8 hr after injection. Data are means ± SEM from n=8 animals; ****p<0.0001, t-test.

To ask whether soma surface-derived biotin-labeled TrkA receptors are transported from ganglia to distal axons *in vivo*, we performed TrkA immunoblotting in streptavidin pulldowns of biotinylated proteins from ganglia and salivary gland lysates. We observed a prominent signal for biotinylated TrkA receptors in the injected superior cervical ganglion, as expected (**Fig. 3B, C**). Notably, we also found biotin-labeled TrkA receptors appearing anterogradely in nerve terminals innervating salivary glands in the injected side (**Fig. 3B**). Biotin-labeled receptors should have originated from soma surfaces since the biotin injected into sympathetic ganglia is membrane-impermeable. We did observe a weaker signal for biotinylated TrkA in contralateral tissues, suggesting that there is a trace amount of biotin that undergoes systemic leak. However, quantification showed a marked enrichment of the biotinylated TrkA signal in injected ganglia (6.5 ± 0.9-fold increase) (**Fig. 3C**) and salivary glands in the injected side (2.2 ± 0.3-fold increase) (**Fig. 3D**) compared to the non-injected side.

To address the specificity of *in vivo* transcytosis of membrane proteins, we also performed immunoblotting for NCAM, a membrane protein abundantly expressed in cell bodies and axons, in streptavidin pulldowns of biotinylated proteins from ganglia and salivary glands. In contrast to TrkA, while biotin-labeled NCAM was enriched in the injected superior cervical ganglion (**Fig. 3B, E**), we barely detected any biotinylated NCAM in pulldowns from salivary glands (**Fig. 3B, F**), suggesting that transcytosis is a selective mechanism to deliver specific membrane proteins, such as TrkA, from soma surfaces to axons.

To visualize TrkA transcytosis *in vivo*, we used *Ntrk1^Flag^*mice in conjunction with local injection of anti-Flag antibodies to label soma surface-TrkA receptors as described above (**Fig. 3A**). 8 hr after injection, superior cervical ganglia and target tissues, iris muscles and salivary glands, were dissected and processed for Flag and Tuj1 immunofluorescence, where Tuj1 is a pan-neuronal marker. Of note, we used anti-mouse Tuj1 antibody to mark neurons instead of the more specific sympathetic marker, Tyrosine Hydroxylase, given that the anti-Flag and anti-TH antibodies are both generated in rabbit. We found that the FLAG immunofluorescence was evident only in the injected ganglion, with no detectable signal in the non-injected side (**Fig. 3G-J**). Further, Flag-TrkA receptors were observed in intracellular puncta within neurons (**Fig. 3K, L**), suggesting internalization of Flag antibody-bound receptors. In assessing target fields, we observed Flag-TrkA immunofluorescence distributed in discrete puncta along the axons innervating the iris muscle (**Fig. 3M-O**) and salivary glands (**Supplementary Fig. 3A-C**) in the injected, but not non-injected, side after 8 hr. The Flag-TrkA signal in nerve terminals was completely abolished by sympathetic axotomy (**Supplementary Fig. 3D-F**), supporting that the appearance of FLAG-antibody labeled TrkA receptors in nerve terminals is due to axonal transport and not leakage of antibodies to distal axons.

Together, these results demonstrate that soma surface-derived TrkA receptors are anterogradely transported to nerve terminals *in vivo*, establishing receptor transcytosis as a physiological mode of axon targeting.

### TrkA transcytosis enhances the formation of presynaptic varicosities *in vitro*

Previously, we reported that soma surface-derived TrkA receptors were localized to axonal growth cones in developing sympathetic neurons ^7^. In this study, we observed that Flag-TrkA receptors were also enriched at discrete swellings along axonal shafts, which were defined by dense accumulation of Tyrosine Hydroxylase (TH) (**Fig. 4A**). TH is the rate limiting enzyme in the biosynthesis of the sympathetic neurotransmitter, Norepinephrine (NE). In post-ganglionic sympathetic neurons, axonal varicosities that appear as bouton-like structures along axon shafts have been characterized as presynaptic compartments that are the sites of neurotransmitter release ^13, 14, 15, 16^. Indeed, we found that the synaptic proteins, synaptophysin-1 and synaptotagmin-1/2, were localized at sites of TH accumulation in axonal varicosities, suggesting that these are presynaptic sites (**Fig. 4B-D**). Live-antibody feeding to monitor the trajectory of soma surface-derived Flag-TrkA receptors, together with immunofluorescence for synaptic proteins, revealed that transcytosed TrkA receptors were enriched at sites of synaptophysin-1 (**Fig. 4 E, F)** or synaptotagmin-1/2 accumulation (**Supplementary Fig. 4A)** along the axon. Further, using EM, we visualized Flag-TrkA gold particles, originally labeled on soma surfaces, appearing at the plasma membrane in axonal varicosities that were defined by their morphology, accumulation of small and large dense-core vesicles, as well as mitochondria (**Supplementary Fig. 4B).** Together, these results suggest that transcytosing TrkA receptors are enriched at presynaptic varicosities in sympathetic axons.

**Figure 4.**
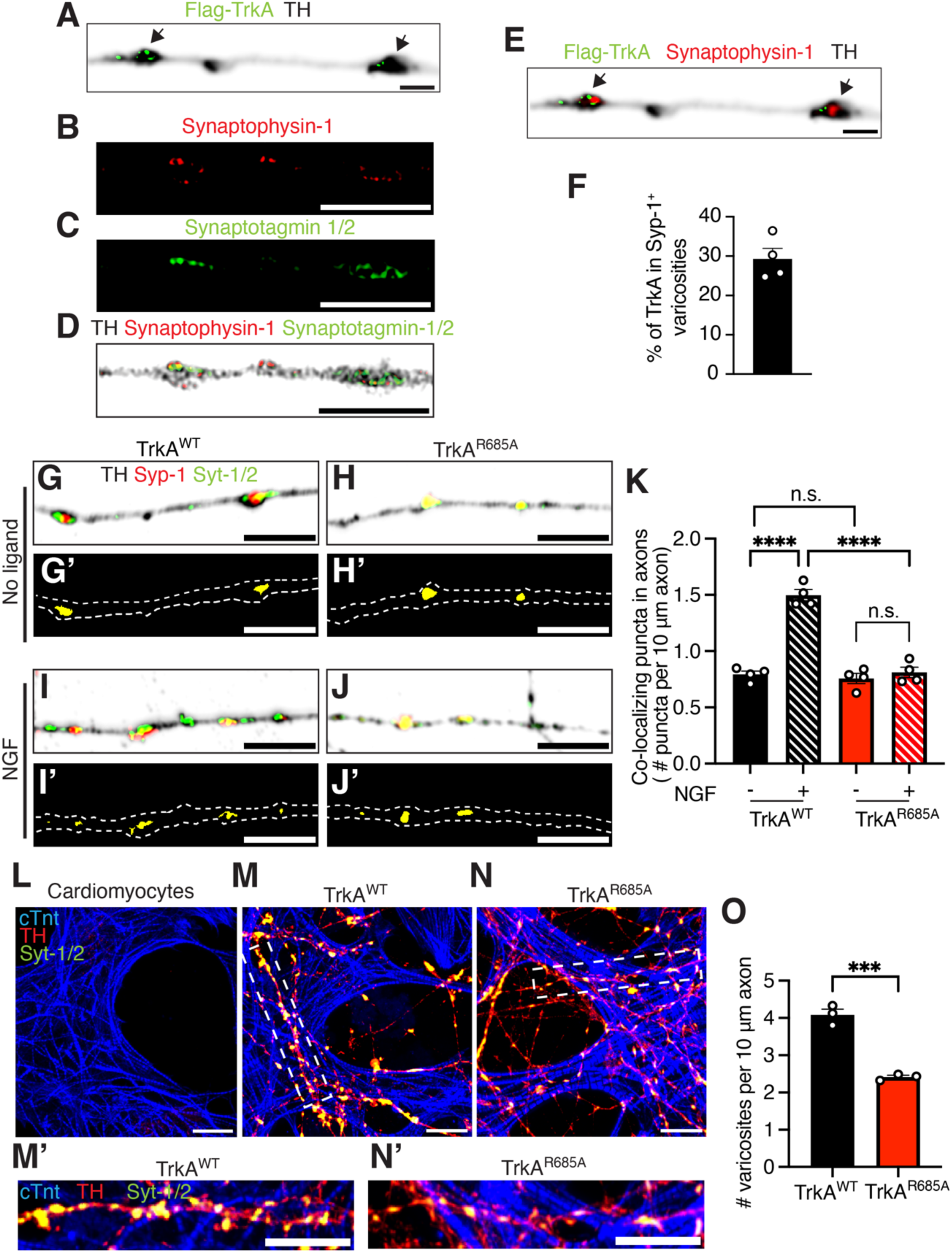
TrkA transcytosis promotes the formation of presynaptic varicosities *in vitro*. (**A**) Soma surface-derived Flag-TrkA puncta accumulate in TH-dense varicosities (arrows) in sympathetic axons. Flag antibody feeding was done in soma + proximal axon compartments in Ntrk1^Flag^ sympathetic neurons grown in microfluidic chambers in the presence of NGF (100 ng/mL). Flag-TrkA (green) and TH (black). Scale bar 2 μm. (**B-D**) Enrichment of synaptophysin-1 (red) and synaptotagmin-1/2 (green) in axonal varicosities. Sympathetic neurons were labeled by TH immunostaining (black). Scale bar 5 μm. (**E**) Soma surface-derived Flag-TrkA (green) accumulate in synaptophysin-1-positive (red) axonal varicosities (arrows). TH labeling is shown in black. Note that the image in (**E**) is the same as that in (**A**) with an additional channel to show synaptophysin-1 immunofluorescence. Scale bar 2 μm. (**F**) Percentage of Flag-TrkA localized in synaptophysin-1 (Syp-1)-positive varicosities relative to total Flag-TrkA puncta in distal axons. Data are means ± SEM from n=4 independent experiments. (**G-J’**) NGF stimulation enhances formation of presynaptic varicosities in TrkA^WT^, but not, TrkA^R685A^ mutant sympathetic neurons. Sympathetic neurons were immunostained with synaptophysin-1 (red), synaptotagmin-1/2 (green) and TH (black). Co-localized synaptophysin-1 and synaptotagmin-1/2 signals to identify presynaptic sites are shown in panels (**G’, H’, I’, and J’**). Scale bar 5 μm. (**K**) Quantification of presynaptic sites in TrkA^WT^ or TrkA^R685A^ sympathetic axons in the presence or absence of NGF (100 ng/mL, 180 min). Presynaptic sites are defined as the number of co-localized synaptophysin-1 and synaptotagmin-1/2 puncta per every 10 μm axon segment. Data are means ± SEM from n=4 independent experiments (total of 80-90 axons); ****p<0.0001, n.s: not significant; two-way ANOVA with Tukey Kramer post-hoc test. (**L-N**) TrkA^R685A^ sympathetic axons make fewer synaptic contacts with cardiomyocytes in co-cultures. Presynaptic sites were identified by co-localization of synaptotagmin-1/2 (green) and TH (red) immunofluorescence in axons at sites of contacts with cardiomyocytes in TrkA^WT^ (**M**) or TrkA^R685A^ (**N**) co-cultures. Cardiomyocytes were labeled by labeled cardiac Troponin t (cTnt, blue). Higher magnification images of insets in panels (**M, N**) are shown in panels (**M’, N’**). Cardiomyocytes cultured alone are shown in (**L**). Scale bar: 10 μm. (**O**) Quantification of the number of varicosities per 10 μm axonal segments in individual axons contacting cardiomyocytes in TrkA^WT^ or TrkA^R685A^ neuron co-cultures. Varicosities were defined by co-localization of synaptotagmin-1/2 and TH in axons. Data are means ± SEM from n=3 independent experiments (total of 25-30 cardiomyocytes); ***p<0.001, t-test.

NGF:TrkA signaling is known to promote the development of presynaptic sites and enhance synaptic transmission in cultured sympathetic neurons ^17, 18, 19^. Given our findings that TrkA receptors are enriched at presynaptic sites after transcytosis, we asked whether receptor transcytosis is required for the formation of presynaptic varicosities in response to NGF stimulation. To disrupt TrkA transcytosis, we used TrkA^R685A^ mice, where TrkA receptor signaling and retrograde trafficking are preserved, but anterograde receptor transcytosis is specifically perturbed ^20^. We established cultures of sympathetic neurons in microfluidic chambers that were harvested either from TrkA^R685A^ mutant or control (TrkA^WT^) littermate mice at postnatal stages (P0-P3). After 1-week, distal axons were stimulated with NGF (100 ng/mL, 180 min) and co-colocalization of synaptophysin-1 and synaptotagmin-1/2 along the axons was measured as a readout of presynaptic sites. We found that NGF treatment significantly increased the number of presynaptic sites in TrkA^WT^ neurons (**Fig. 4G, G’, I, I’, K**). However, the NGF-induced increase in number of presynaptic sites was abrogated in TrkA^R685A^ mutant neurons (**Fig. 4H, H’, J, J’, K**).

As an additional measure to assess the requirement for TrkA transcytosis in presynaptic assembly, we resorted to co-cultures of sympathetic neurons with cardiomyocytes, a well-established *in vitro* system to study the formation of presynaptic contacts and synaptic communication between sympathetic neurons and target effector cells ^17, 18^. In co-cultures, sympathetic neurons establish functional synaptic contacts with cardiomyocytes and influence their spontaneous beat rate *in vitro* ^17, 18, 21^. We established co-cultures of either TrkA^WT^ or TrkA^R685A^ mutant sympathetic neurons with wild-type cardiomyocytes in microfluidic chambers, where the cardiomyocytes were plated in axonal compartments (**Supplementary Fig. 4C)**. Although cardiomyocytes express NGF, the culture media was supplemented with NGF (100 ng/mL) to ensure neuronal viability. We observed that both TrkA^WT^ and TrkA^R685A^ sympathetic neurons make synaptic connections with cardiomyocytes, where presynaptic varicosities along axons were identified by their bouton-like morphology and co-localization of synaptotagmin-1/2 and TH (**Fig. 4L-N’**). Cardiomyocytes were visualized by cardiac troponin T (cTnt) immunostaining. However, there was a significant decrease in the number of presynaptic varicosities in TrkA^R685A^ mutant sympathetic processes in co-cultures (**Fig. 4N, N’, O**), compared to TrkA^WT^ neurons (**Fig. 4M, M’, O**), despite similar extent of axon outgrowth into distal compartments (**Supplementary Fig. 4D-F).**

Together, these results support that transcytosed TrkA receptors are enriched at presynaptic varicosities along sympathetic axons, and that TrkA transcytosis promotes the formation of presynaptic sites.

### TrkA transcytosis enhances synaptic transmission in neuron-cardiomyocyte co-cultures

We next asked whether TrkA transcytosis is critical for sympathetic synaptic transmission. The majority of sympathetic neurons, including those innervating the heart, are noradrenergic, releasing Norepinephrine (NE) as their primary neurotransmitter. To monitor NE release, we employed a genetically encoded NE biosensor, G-protein-coupled receptor (GPCR)-activation-based (GRAB) NE sensor called GRAB_NE2h_, in which NE binding induces a conformational change in the α2 adrenergic receptor (α2AR) to drive a fluorescence change in circular permutated EGFP (cpEGFP) ^22^. Expression of GRAB_NE2h_ in cardiomyocytes using an adenoviral vector resulted in a robust increase after NE (100 µM) application (**Fig. 5A, B**), confirming biosensor functionality. To assess synaptic communication between sympathetic neurons and cardiomyocytes, we established co-cultures of TrkA^WT^ or TrkA^R685A^ mutant sympathetic neurons with wild-type cardiomyocytes in microfluidic chambers (**Fig. 5C**). After 5 days in co-cultures to allow axon outgrowth into distal compartments, cardiomyocytes were infected with the adenoviral construct co-expressing GRAB_NE2h_ and mCherry. Live-cell imaging to monitor NE signaling was performed in GRAB_NE2h_-expressing cardiomyocytes that were identified by mCherry expression (**Supplementary Fig. 5A, B)**. Further, we specifically imaged cardiomyocytes at sites of contact with sympathetic axons, where axons were live-labeled by anterogradely transported Alexa Fluor 647-conjugated cholera toxin subunit B that had been added to cell bodies (**Fig. 5C and Supplementary Fig. 5A, B**). To trigger NE release, nicotine (10 µM) was added specifically to the cell body compartments (**Fig. 5C**), where nicotine acting on nicotinic Acetylcholine receptors expressed in neuronal cell bodies, depolarizes sympathetic neurons and stimulates NE-mediated neurotransmission ^23^. In co-cultures with TrkA^WT^ or TrkA^R685A^ sympathetic neurons, we observed a basal level of GRAB_NE2h_ fluorescence in cardiomyocytes indicative of spontaneous neuronal activity (**Fig. 5D, F**). Nicotine application to cell bodies resulted in a pronounced increase in biosensor fluorescence in cardiomyocytes co-cultured with TrkA^WT^ sympathetic neurons compared to TrkA^R685A^ mutant neurons (**Fig. 5E, G**). Quantification revealed a significant (4.2-fold) increase in ΔF/F0 in GRAB_NE2h_ fluorescence in response to nicotine in cardiomyocytes co-cultured with wild-type compared to mutant sympathetic neurons (**Fig. 5H**). These results suggest that disruption of TrkA transcytosis impairs synaptic transmission.

**Figure 5.**
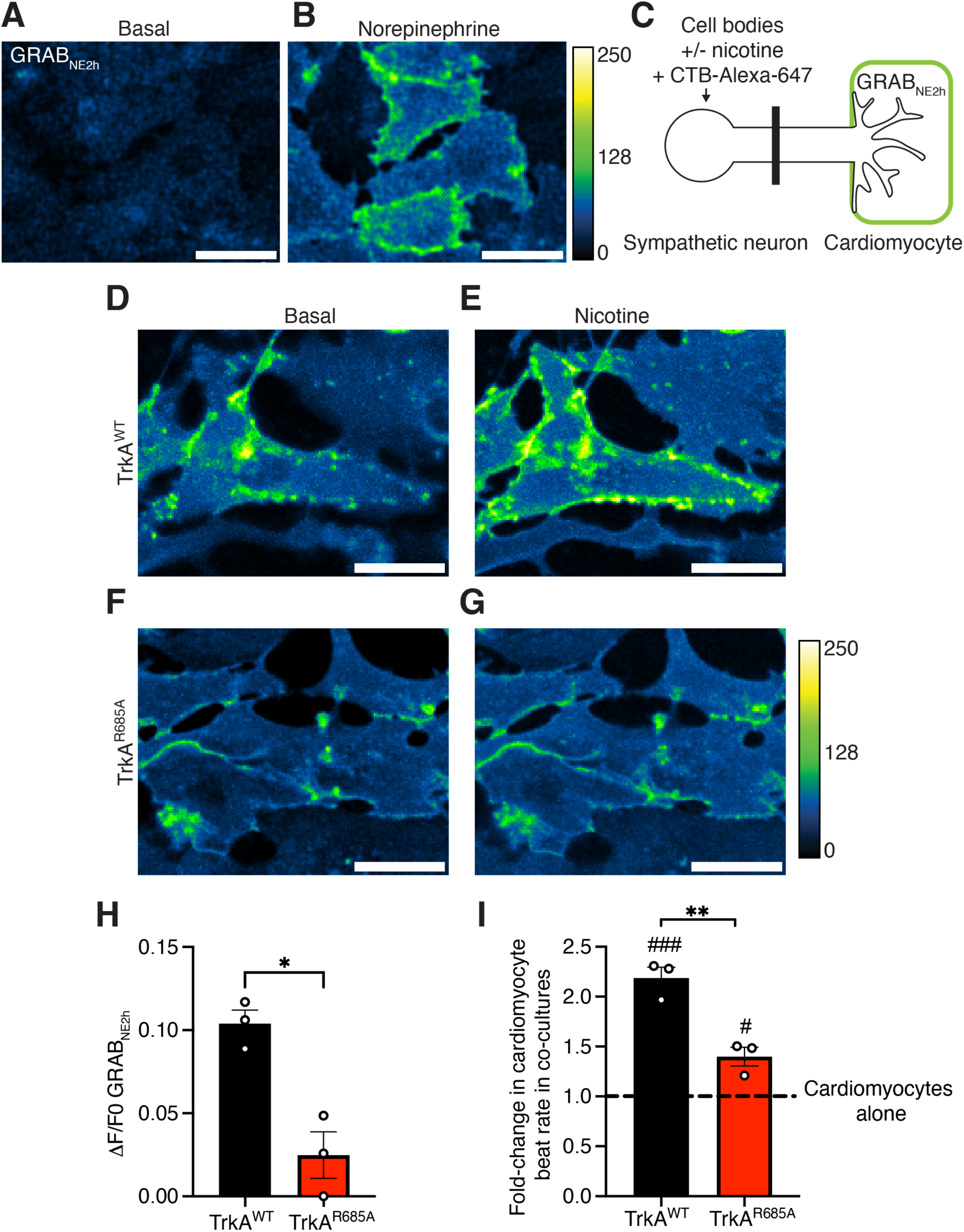
TrkA transcytosis enhances synaptic transmission in neuron-cardiomyocyte co-cultures. (**A-B**) Cardiomyocytes expressing the norepinephrine (NE) biosensor (GRAB_NE2h_) show a robust increase in fluorescence in response to NE (100 μM) application. Calibration bar for fluorescence intensity is shown to the right of the panels. Scale bar 20 μm. (**C**) Schematic of sympathetic neuron-cardiomyocyte co-cultures in microfluidic chambers to assess NE release in response to nicotine (10 μM) added specifically to soma + proximal axon compartments. Cardiomyocytes were plated in distal axon compartments and infected with an adenovirus expressing CMV-GRAB_NE2h_-EF1a-mCherry. Alexa-647-Cholera Toxin Subunit B (CTB-Alexa-647, 1 µg/mL) was added specifically to soma + proximal axon compartments for anterograde labeling to visualize distal axons contacting cardiomyocytes. (**D-G**) Enhanced GRAB_NE2h_ fluorescence in cardiomyocytes cultured with TrkA^WT^ sympathetic neurons compared to TrkA^R685A^ mutant neurons in response to nicotine stimulation of neuronal soma. Calibration bar for fluorescence intensity is indicated at the bottom of the panels. Scale bar 20 μm. (**H**) Quantification of cardiomyocyte-associated GRAB_NE2h_ fluorescence (ι1F/F0) in response to nicotine stimulation of co-cultured TrkA^WT^ or TrkA^R685A^ sympathetic neurons. Data are means ± SEM from n=3 independent experiments (total of 20 cardiomyocytes); *p<0.05, t-test. (**I**) Cardiomyocyte beat rate is significantly enhanced by co-culturing with TrkA^WT^ compared to TrkA^R685A^ sympathetic neurons. Values are represented as fold-change in cardiomyocyte beat rate in TrkA^WT^ or TrkA^R685A^ co-cultures, compared cardiomyocyte cultures alone (dashed line). Data are means ± SEM from n=3 independent experiments; **p<0.01 for TrkA^WT^ compared to TrkA^R685A^, ### p<0.001 and # p<0.05 for TrkA^WT^ and TrkA^R685A^ compared to cardiomyocytes alone, respectively; one-way ANOVA and Tukey-Kramer test.

As an additional measure of functional connections between sympathetic neurons and cardiomyocytes, we measured the spontaneous beat rate of cardiomyocytes in culture, which is enhanced in the presence of sympathetic neurons ^18, 21^. We found that cardiomyocyte beat rate is significantly enhanced by co-culturing with TrkA^WT^ compared to TrkA^R685A^ sympathetic neurons (**Fig. 5I, Supplementary movies 3-5**).

Together, these results provide evidence that TrkA receptor transcytosis is critical for synaptic transmission in sympathetic neurons.

### TrkA transcytosis contributes to presynaptic specialization of sympathetic nerve terminals *in vivo*

We next asked if TrkA transcytosis promotes the formation of presynaptic sites in sympathetic nerve terminals *in vivo*. Compared to the abundant knowledge about synapse formation in the central nervous system (CNS) and neuromuscular junctions, relatively little is known about how sympathetic axons make contacts with peripheral targets. Limited insight has come from classical ultrastructural studies, where neurotransmitter release is thought to occur from varicosities that contain clusters of small clear and large dense-core vesicles distributed along axonal shafts ^13, 14, 24^, similar to observations in sympathetic neurons in culture. However, relatively little is known about the assembly of presynaptic varicosities and/or the molecular/cellular underpinnings. To assess presynaptic development *in vivo*, we turned to the iris dilator muscle that is densely innervated by sympathetic nerves, causing muscle contraction through noradrenergic neurotransmission to drive pupil dilation. Previous studies have established that sympathetic neurotransmission is proposed to occur from presynaptic varicosities that are closely apposed to myoepithelial cells, pigment cells (melanocytes) and vascular smooth muscle cells ^25, 26, 27^.

To visualize the formation of presynaptic varicosities in the iris across development, we employed TH and synaptophysin-1 immunohistochemistry in flat-mount preparations of eye tissue, that specifically include the cornea and iris, at different postnatal stages in mice. As developmental time-points, we chose postnatal day 1 (P1), when iris sympathetic innervation is just beginning, postnatal day 15 (P15), when the sympathetic innervation gains functionality based on muscle contractility, and postnatal day 30 (P30), when sympathetic innervation of iris is complete ^26, 28^. At P1, we observed a sparse plexus of sympathetic fibers in the iris, where the axons were relatively smooth with low levels of synaptophysin-1 expression (**Fig. 6A, A’, D**). By P15, there was a pronounced increase in sympathetic innervation with an increase in the number of presynaptic varicosities, identified by their morphology, as well as accumulation of synaptophysin-1 and TH (**Fig. 6B, B’ and D**). By P30, most of the sympathetic axons in the iris had a varicose, beads-on-a string-like appearance with a significant increase in the number and volume of the presynaptic varicosities compared to earlier stages (**Fig. 6C, C’, D, E**). These results suggest that presynaptic varicosities in sympathetic neurons are formed during the first 3-4 postnatal weeks in mice.

**Figure 6.**
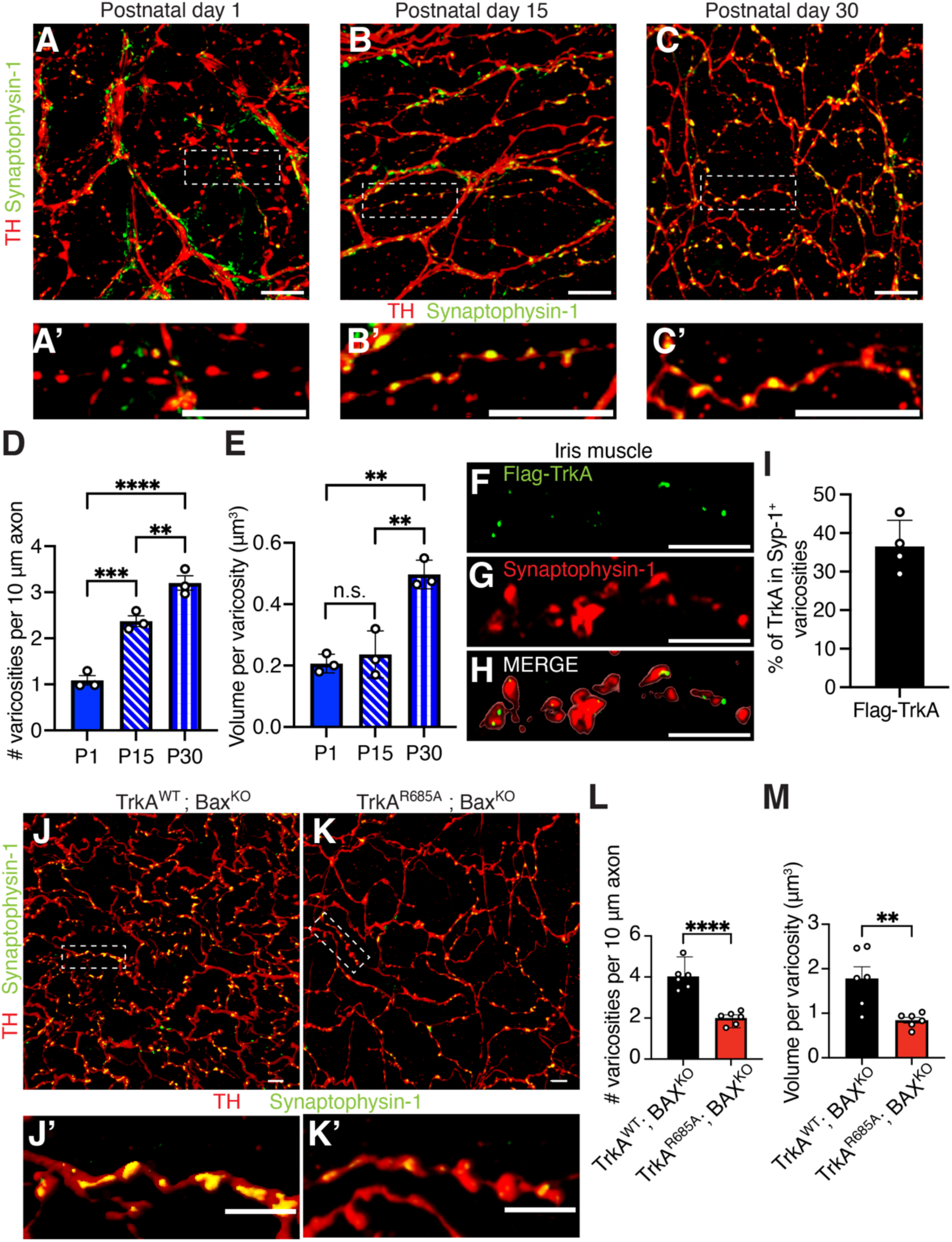
TrkA transcytosis promotes the formation of presynaptic varicosities *in vivo*. (**A-C**) Postnatal development of presynaptic varicosities in sympathetic innervation of the iris revealed by synaptophysin-1 (green) and TH (red) immunofluorescence. Sympathetic nerves in the iris dilator muscle were assessed at postnatal days 1, 15, and 30. Higher magnification images of insets in (**A, B, C**) are shown in panels (**A’, B’, C’**). Scale: 10 μm. (**D-E**) Developmental increase in the number and size of varicosities. Number of varicosities were quantified per 10 μm axon segments (**D**) as well as volume of varicosities (**E**) in individual axons in wild-type mice. Data are means ± SEM from n=3 animals for each postnatal stage; **p<0.01, ***p<0.001, ****p<0.0001, n.s: not significant, one-way ANOVA and Tukey-Kramer test. (**F-H**) Transcytosed Flag-TrkA (green) appear in synaptophysin-1-positive (red) varicosities in sympathetic axons innervating the iris dilator muscle. FLAG and synaptophysin-1 immunofluorescence was done 8 hr after injection of Flag-antibodies into ganglia (SCG) in P30 Ntrk1^Flag^ mice. Scale: 5 μm. (**I**) Quantification of Flag-TrkA puncta in synaptophysin-1-positive varicosities as a percentage of total Flag-TrkA signal in distal axons. Data are means ± SEM from n=4 mice. (**J-K’**) TrkA^R685A^:Bax^KO^ mice have reduced number as well as aberrant morphology of presynaptic varicosities compared to TrkA^WT^:Bax^KO^ animals. Presynaptic varicosities were visualized by immunostaining for synaptophysin-1 (green) and TH (red). Higher magnification images of insets in panels (**J, K**) are shown in panels **(J’, K’**). Scale 5 μm. (**L, M**) Quantification of varicosity number (per 10 μm axon) and volume in individual axons. Data are means ± SEM from n=6 mice per genotype; **p<0.01, ****p<0.0001, t-test.

We next asked whether TrkA is targeted to presynaptic varicosities *in vivo*. Thus, we injected anti-Flag antibodies to the superior cervical ganglia in *Ntrk1^Flag^* mice at P30, when presynaptic terminals are fully established and functional ^26, 28^. After performing whole mount TH and synaptophysin-1 immunofluorescence, we visualized Flag-TrkA particles, originating from ganglia, localized in synaptophysin-1-positive varicosities in sympathetic nerve terminals innervating the iris (**Fig. 6F-H**). Quantification revealed that 36.5 ± 3.4% of Flag-TrkA particles were localized in presynaptic varicosities in the iris (**Fig. 6I**). Similarly, we also observed soma surface-derived FLAG-TrkA particles appearing in presynaptic varicosities in the salivary glands, (**Supplementary Fig. 6A-C),** with 39.3 ± 5.9% of Flag-TrkA particles localized in presynaptic varicosities (**Supplementary Fig. 6D)**. These results indicate that transcytosed TrkA receptors are localized to presynaptic varicosities *in vivo*, consistent with the results in cultured sympathetic neurons.

Lastly, we addressed whether TrkA transcytosis contributes to the formation of presynaptic varicosities *in vivo*. We previously reported that disruption of TrkA transcytosis in TrkA^R685A^ mice results in decreased axonal TrkA levels *in vivo*, leading to the developmental loss of sympathetic neurons (∼40% decrease by birth) and reduced innervation of target tissues ^20^. To assess presynaptic varicosities without the complication of neuronal apoptosis in TrkA^R685A^ mice, we concomitantly deleted *Bax*, an essential gene for neuronal apoptosis ^29^, to generate TrkA^R685A^:Bax^KO^ double-mutant mice. As controls, litter-mate TrkA^WT^:Bax^KO^ mice were used. Wholemount TH and synaptophysin-1 immunostaining revealed a marked decrease in the number of presynaptic sites, assessed by synaptophysin-1 enrichment in varicosities, in TrkA^R685A^:Bax^KO^ mice compared to control animals (**Fig. 6J, J’, K, K’ and L**). Quantification revealed that there were 4.0 ± 0.3 presynaptic varicosities per 10 µm axon segment in control animals, compared to 2.0 ± 0.1 in the mutants (**Fig. 6L**). Further, presynaptic varicosities were much smaller in TrkA^R685A^:Bax^KO^ axons, with a 53% decrease in volume compared to control axons (**Fig. 6M**). Of note, TrkA^R685A^:Bax^KO^ mice had reduced axon innervation relative to control mice (**Fig. 6J, K**), despite the rescue of neuronal survival, indicating that axon innervation is still affected by the disruption of receptor transcytosis. However, since we assessed presynaptic varicosities in individual axons, the attenuated number and morphology of axonal varicosities indicate impaired formation of presynaptic sites that is independent of the density of innervation.

Together, these results suggest that TrkA receptor transcytosis is a critical determinant in the establishment of presynaptic sites in sympathetic axon terminals *in vivo*.

## Discussion

Neuronal responsiveness to target-derived factors requires the precise axonal targeting of new receptors, synthesized in cell bodies, to their functional sites in axons. In this study, we provide mechanistic and functional insight into an unconventional mode of ligand-triggered transcytosis of TrkA receptors from soma surfaces to axons in sympathetic neurons. We define the dynamic behavior of transcytosing TrkA receptors in axons and show that soma-surface receptors undergo sorting from being housed in multivesicular bodies and endosomes in proximal axons to a primarily endosomal localization in distal axons. We demonstrate that TrkA transcytosis occurs *in vivo*, that transcytosed TrkA receptors are enriched in presynaptic axonal varicosities, and that receptor transcytosis enhances presynaptic specialization during postnatal development in mice. Using a co-culture system of sympathetic neurons and cardiomyocytes, in combination with a NE biosensor, we show that TrkA transcytosis is critical for noradrenergic synaptic communication between sympathetic neurons and target effector cells. Together, these findings provide fundamental insight into a receptor trafficking pathway that shapes sympathetic nervous system connectivity and function.

Retrograde trafficking of axon-derived TrkA receptors has been the focus of intense research over several decades, with the thorough characterization of transport dynamics and trafficking itinerary ^3, 4, 30^. In agreement with previous studies ^11, 12^, we found that axon-derived TrkA receptors move solely in a retrograde direction in a saltatory manner with few intermittent pauses in axons. In contrast, soma surface-derived TrkA receptors move in a markedly different manner with distinct behaviors in proximal versus distal axons. Live-imaging revealed that NGF stimulation of distal axons triggers the rapid anterograde movement of ∼25% of soma surface-derived TrkA receptors in proximal axons, whereas the majority of soma surface-derived receptors appeared to be stationary. Stationary receptors may reflect those that are still resident on the plasma membrane or targeted for recycling and/or degradation after internalization. This is consistent with our ultrastructural analyses, where soma surface-receptors were observed at the plasma membrane, lysosomes, and multivesicular bodies. Intriguingly, in distal axons, we found bi-directional movement of soma surface-derived TrkA receptors, with approximately 25% of receptors undergoing retrograde movement. It is possible that this population reflects soma surface-derived receptors that were inserted on the plasma membrane and then undergo retrograde trafficking to propagate NGF signaling to cell bodies. However, their retrograde speed (1.18 ± 0.08 µm/s) was markedly slower than that of axon-derived TrkA receptors (1.59 ± 0.10 µm/s) suggesting that these are distinct populations. Since all retrogradely transported organelles use the one dynein motor, the different speeds may reflect the use of different adaptor proteins. Alternatively, it is possible that retrogradely moving soma surface-derived receptors were never inserted on the plasma membrane, but that they switch directions within distal axons because of dynamic contacts with cytoskeletal elements, a dense actin meshwork for example ^31^, or other organelles such as axonal ER ^32^.

Our findings also suggest a sorting of the organelles carrying transcytosed TrkA receptors in the proximal axons. Using EM, we found a similar distribution of soma surface-derived TrkA receptors in multivesicular bodies (41.6%) and endosomes (34.2%), in proximal axons, which shifted to receptors being predominantly localized in endosomes (66%) in distal axons. Previously, we reported that soma surface-labeled Trk receptors co-localized with Rab11a, a marker for long-distance recycling endosomes, in sympathetic neurons ^7^. Further, inhibition of Rab11a activity disrupted TrkA transcytosis ^7^. These results suggest that the single membrane vesicles carrying soma surface-derived TrkA receptors in axons are likely Rab11a-positive recycling endosomes. Multivesicular bodies typically fuse with lysosomes or with the plasma membrane to generate exosomes ^33, 34, 35^. It is possible that soma surface-derived TrkA receptors found in multivesicular bodies reflect receptors that are either targeted for lysosomal degradation or perhaps secreted in exosomes from the plasma membrane. Intriguingly, multivesicular bodies carrying axon-derived TrkA receptors have been reported to mature into single-membrane TrkA-positive vesicles upon arriving in cell bodies ^36^. Thus, an alternative fate of multivesicular bodies carrying soma surface derived TrkA receptors in proximal axons is that they mature into single-membrane TrkA-positive vesicles that then travel to distal axons.

Based on studies in neuronal cultures, transcytosis has been proposed as a mechanism for axonal delivery for a limited number of membrane proteins, including NgCAM, a cell adhesion molecule important for axon guidance ^37^, β-1-integrin receptors known to mediate growth cone motility and neurite outgrowth^38^, and the type 1 cannabinoid receptor (CB1R), an abundant G-protein coupled receptor implicated in synaptic plasticity ^39^. Further, impaired transcytosis of amyloid precursor protein and low-density lipoprotein receptors has been observed in human iPSC-derived neurons with mutations in familial Alzheimer’s Disease-associated genes ^40^. In this study, we established an *in vivo* strategy to label soma surface TrkA receptors residing in sympathetic ganglia that allowed us to demonstrate that transcytosis is a physiological mode of receptor delivery to nerve terminals. Our findings provide the opportunity to assess whether additional membrane proteins that are critical for axon growth, neuronal survival, and synaptic functions undergo transcytosis *in vivo*. It will be of interest in the future to combine *in vivo* cell-surface biotin labeling of membrane proteins in neuronal cell bodies followed by streptavidin pull-downs of biotinylated proteins that have been anterogradely transported to axons with mass spectrometry to globally identify membrane proteins that undergo transcytosis in neurons.

Using TrkA^R685A^ knock-in mice, where a point mutation in TrkA abolishes receptor transcytosis ^20^, we reveal a critical role for TrkA transcytosis in the formation of functional presynaptic sites in sympathetic neurons. The formation of synaptic contacts between sympathetic neurons and their peripheral targets is a poorly understood process. Compared to the classical chemical synapses in the CNS, post-ganglionic sympathetic axons have large *en passant* boutons or varicosities that are the reported sites of neurotransmitter release, with no post-synaptic specializations ^13, 14, 41^. The molecular mechanisms governing the assembly, maturation, and function of these presynaptic sites remain largely unknown. However, NGF signaling is known to acutely potentiate synaptic communication between sympathetic neurons and cardiomyocytes, as well as exert a long-term effect on increasing the number and/or strength of synapses, in co-cultures ^18^. We found that formation of presynaptic sites *in vivo* occurs over the first 3-4 weeks after birth in mice, as evidenced by an increase in the number and size of the varicosities as well as accumulation of synaptic proteins and TH in varicosities. Disruption of TrkA transcytosis resulted in fewer and smaller presynaptic varicosities in TrkA^R685A^ mutant neurons *in vitro* and *in vivo*, as well as attenuated NE-mediated neurotransmission in co-cultures. Given that axonal TrkA protein levels are reduced by ∼70% in TrkA^R685A^ mice ^20^, the defects in presynaptic site formation and function in mutants could simply reflect reduced receptor availability in axons, which would compromise neuronal responsiveness to target-derived NGF. Notably, NGF signaling is required for the transcriptional regulation of several genes encoding for synaptic proteins, including α-synuclein, synaptotagmin-9, synapsin II, and neurexin 1, in sympathetic neurons ^42^. However, our observations that transcytosed TrkA receptors are localized at presynaptic varicosities together with synaptic proteins suggest that receptor transcytosis could play a more direct and instructive role in the formation of synaptic connections, potentially by actively recruiting presynaptic machinery to initiate the *de novo* formation of presynaptic sites or to stabilize pre-existing presynaptic protein assemblies.

Our study addresses several key questions about an under-appreciated pathway for long-distance axonal targeting of TrkA receptors in sympathetic neurons, including insight into the cell biology of receptor transcytosis and its physiological relevance in shaping sympathetic nervous system connectivity and synaptic function. Since basic principles of axonal delivery are likely to be shared amongst neurons, transcytosis might be a more general mechanism than currently appreciated for targeting of trophic and guidance receptors, adhesion and synaptic proteins, as well as ion channels in other neuronal populations. Uncovering mechanisms of axon delivery have implications that extend beyond the healthy nervous system to understanding cell biological pathways that contribute to nerve repair after injury or neurodegeneration, since the correct complement of membrane proteins must be accurately targeted to regenerating axons to ensure functional recovery.

## Materials and Methods

### Key Resources Table

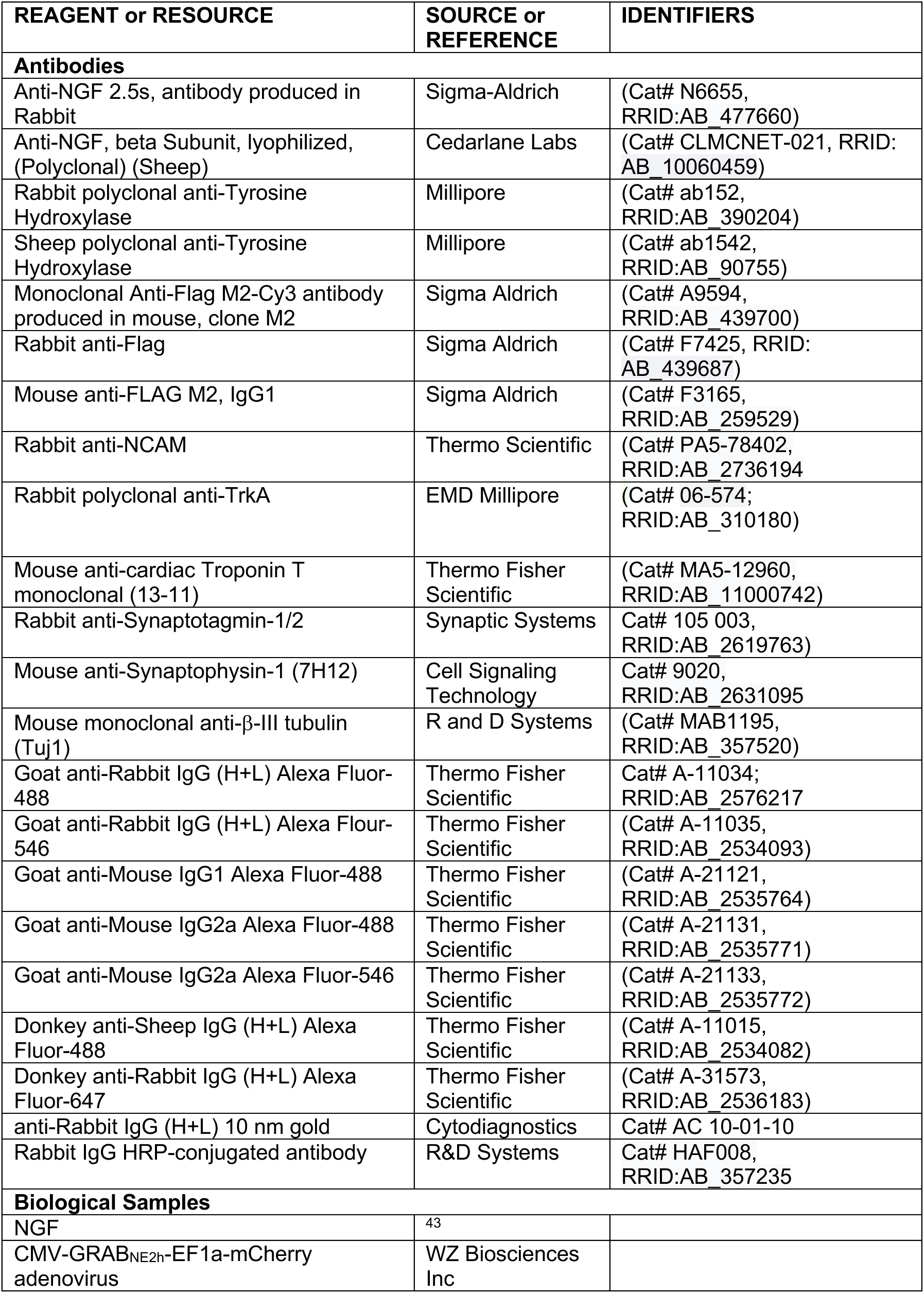
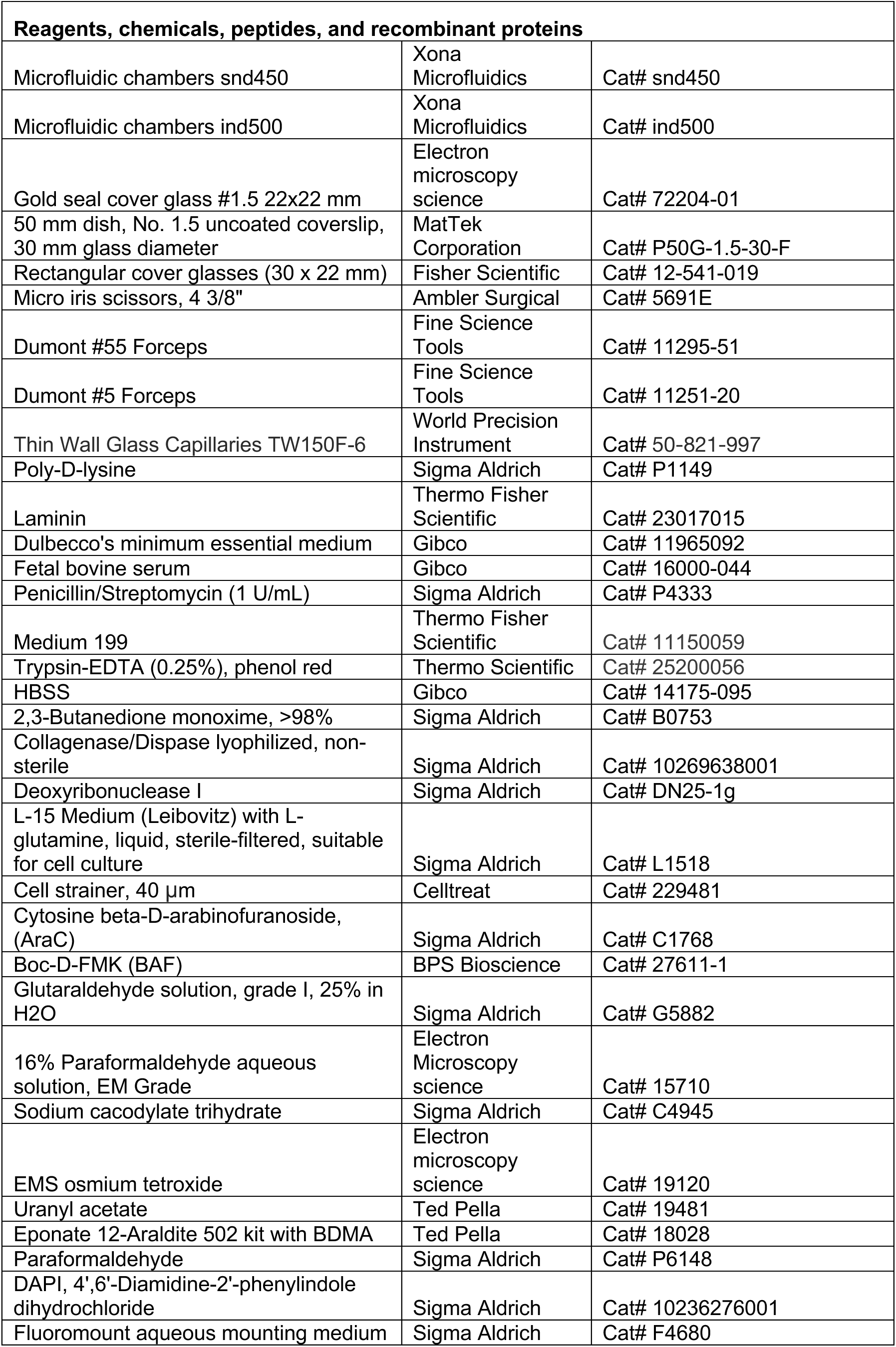
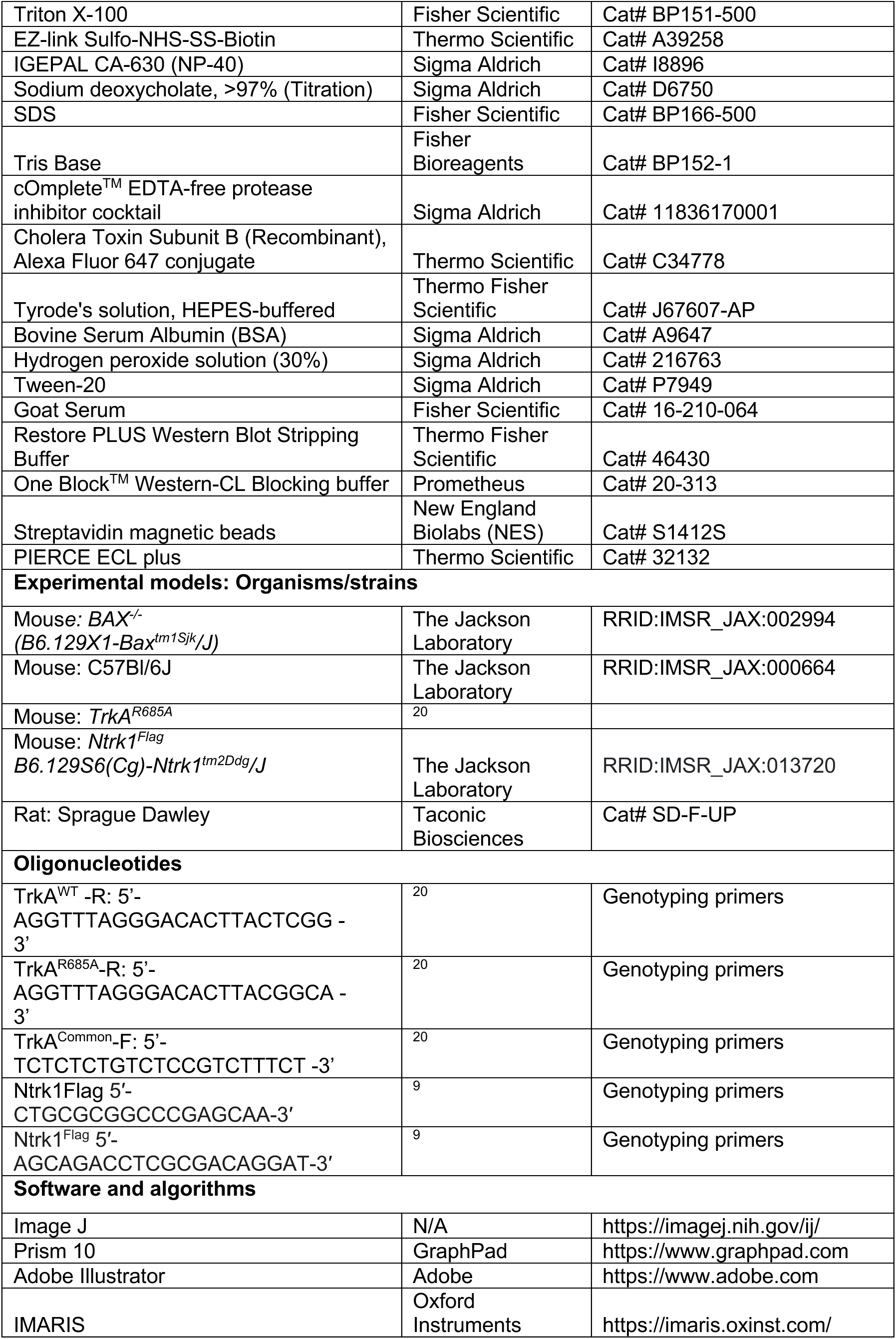

## Methods

### Animals

All animal care and experimental procedures were conducted in accordance with the Johns Hopkins University Animal Care and Use Committee (ACUC) and NIH guidelines. All efforts were made to minimize the pain and number of animals used. Animals were group housed in a standard 12:12 light-dark cycle, with excess water and food *ad libitum*. Pregnant untimed Sprague Dawley rats were purchased from Taconic Biosciences. The ages of mice and rats are indicated in the Figure Legends and/or Methods. Both male and female mice and rats were used for the analyses. The following mouse lines were used in this study: control C57BL/6J mice, TrkA^R685A^ knock-in mice that were previously generated in our laboratory ^20^; Ntrk1^Flag^ knock-in mice (*B6.129S6(Cg)-Ntrk1^tm2Ddg^/J*, JAX:013720) and Bax^+/−^ mice (*B6.129X1-Bax^tm1Sjk^*/J, JAX #002994) were purchased from the Jackson Laboratory.

### Sympathetic neuron cultures and co-cultures with cardiomyocytes

Superior cervical ganglia (SCGs) were harvested from mice (Ntrk1^Flag^, TrkA^WT^, or TrkA^R685A^ mice) at postnatal stages P0-P3. Neurons were enzymatically dissociated and cultured on poly-D-lysine/laminin coated glass coverslips mounted in microfluidic chambers. Neurons were cultured in high-glucose DMEM media supplemented with 10% FBS, 1 U/mL penicillin/streptomycin, 100 ng/mL NGF, and 1 μM AraC to prevent proliferative cell growth as described previously ^44^, for 5-7 days *in vitro* to allow axon growth into distal compartments.

For neuron-cardiomyocyte co-cultures, neonatal mouse cardiomyocytes were isolated and prepared according to a protocol adapted from ^45^. Briefly, hearts were dissected from TrkA^WT^ mice at P0-P3, the ventricles were minced using forceps in Trypsin 0.25% in Hanks’ Balanced Salt Solution (HBSS) supplemented with 2,3-Butanedione monoxime (20 mM) and incubated at 37°C for 60 min. Enzymatic digestion was done using Collagenase/Dispase (0.25 mg/mL) dissolved in L-15 media supplemented with DNAse (0.6 mg/mL) and 2,3-Butanedione monoxime (20 mM). After incubation for 30 min at 37 °C, digestion mix was strained through a 40 μm cell strainer and centrifuged for 5 min, 200g, at room temperature. The pellet was resuspended in co-culture media (DMEM supplemented with 15% FBS, penicillin/streptomycin at 1 U/mL, and 20% M-199) and plated in an uncoated 6 cm cell culture dish to remove fibroblasts and endothelial cells, which adhere to the uncoated culture dish. After 60-90 min, supernatant containing non-adherent cardiomyocytes was centrifuged for 5 min at 200g, re-suspended in co-culture media supplemented with NGF (100 ng/mL) and plated in axonal compartments in microfluidic chambers. Co-cultures were maintained for 5-7 days *in vitro*.

### Live imaging

Sympathetic neurons isolated from Ntrk1^Flag^ mice (P0-P3) were grown in microfluidic chambers in culture media containing NGF (100 ng/mL) for 7 days *in vitro* until axons projected into the distal compartments. Neurons were then deprived of NGF by incubating in high-glucose DMEM containing 1% FBS, anti-NGF (1 µg/mL), and BAF (50 μM), a broad-spectrum caspase inhibitor to prevent apoptosis, for 24 hr. Surface Flag-TrkA receptors in soma + proximal axon compartments were live-labeled with monoclonal anti-Flag M2-Cy3 antibodies (1:200) in Phosphate Buffered Saline (PBS) for 30 min at 4°C. Excess antibody was washed off with PBS, followed by stimulation with NGF (100 ng/mL) added specifically to distal axon compartments for 20-180 min at 37°C. Anti-NGF (1:500) was added to soma + proximal axon compartments. Neurons were imaged in CO_2_-buffered 37°C stage-top chamber mounted on a LSM 980 Zeiss confocal microscope equipped with a spectral detector. Images were acquired at two frames per second (512 x 512 pixels) using a 63X oil immersion objective (1.40 NA). Time-lapse images were imported to Image J (NIH) and individual particles were tracked manually and analyzed by the Multi Kymograph plug-in. Distance traveled by Flag-TrkA particles between two paused points was used to determine instantaneous speed and directionality. Data were obtained from three independent experiments with imaging of at least 5 different chambers per condition.

### Electron microscopy

Sympathetic neurons isolated from Ntrk1^Flag^ mice (P0-P3) were cultured in microfluidic chambers as described above. After axons had projected into distal compartments, NGF was withdrawn from culture media for 24 hr. Surface Flag-TrkA receptors in soma + plus proximal axon compartments were live-labeled with Rabbit anti-Flag antibody (1:200) pre-conjugated to anti-Rabbit IgG (H+L)-10 nm gold secondary antibody in PBS for 30 min at 4 °C. Excess antibody was washed off with PBS, followed by stimulation with NGF (100 ng/mL) added to distal axon compartments for 180 min. Microfluidic devices were then removed, neurons washed in PBS, and fixed in 2.5% glutaraldehyde plus 4% PFA in 0.1 M cacodylate buffer for 60 min at room temperature. Neurons were post-fixed with 1% osmium tetroxide, stained with 1% uranyl acetate to enhance membrane contrast, and dehydrated in a series of incubations with ethanol (30%, 50%, 70% 90% and 100%) for 30 min each, followed by embedding in EPON resin. The next day, coverslips were removed and thin sections, 60 to 90 nm, were cut with a diamond knife on a Leica UCT ultramicrotome and mounted on Formvar copper slot grids. Grids were stained with 2% uranyl acetate followed by lead citrate. Images were captured using a Hitachi 7600 transmission electron microscope with a Dual AMT CCD camera at 50,000-80,000X. Over 100 samples were prepared and analyzed from soma + proximal axon as well as distal axon compartments. Distribution of Flag-TrkA-10 nm gold particles in proximal and distal axons was quantified from multiple randomly selected samples from three independent experiments.

### Live antibody feeding and immunostaining

For co-localization of transcytosed TrkA with synaptic proteins, Flag antibody feeding was performed to label soma surface receptors with mouse or rabbit anti-Flag antibodies (1:200) in PBS for 30 min at 4°C. After NGF (100 ng/mL) stimulation of distal axons for 180 min, neurons were fixed with 4% PFA in PBS for 15 min, incubated in blocking buffer (5% BSA, 0.3% Triton X-100 in PBS) for 1 hr at room temperature, and then incubated with primary antibodies, which include mouse anti-synaptophysin-1 (1:1000), rabbit anti-synaptotagmin-1/2 (1:500), or sheep anti-TH antibody (1:500), in blocking buffer overnight at 4°C. Following PBS washes, neurons were incubated with anti-rabbit Alexa-594, anti-mouse Alexa-647, or anti-sheep-488 secondary antibodies (1:500) and DAPI (0.3 µM) in blocking buffer. Samples were washed 3 times with PBS and mounted in Fluoromount aqueous mounting medium. Images were obtained by a Zeiss LSM 980 confocal scanning microscope equipped with Airy Scan module. Images were captured in ∼1-2 µm z-stacks at intervals of 0.15 µm using a 63X objective (1.4 NA) with 2.5x digital zoom with optimal settings for Airy Scan mode. The same acquisition settings were applied to all images taken from a single experiment. Images were processed and analyzed by Image J. Each channel for each image was thresholded. Number of Flag-TrkA puncta localized in varicosities positive for synaptophysin-1 or synaptotagmin-1/2 per 10 µm axon length was quantified and expressed relative to the total Flag-TrkA puncta in distal axons. Puncta were counted using particle analysis function in Image J. In some experiments, the number of co-localizing puncta for synaptotagmin-1/2 and synaptophysin-1 was quantified per 10 µm axon length. In all experiments, TH immunostaining was used to visualize axons. Co-localization analysis was done using Image J.

In sympathetic neuron-cardiomyocyte co-cultures, neurons and cardiomyocytes were labeled by immunostaining using sheep anti-TH (1:500) and rabbit anti-synaptotagmin-1/2 (1:500) for neurons, and a mouse anti-cardiac troponin T (cTnt) (1:1000) for cardiomyocytes. After overnight incubation with primary antibodies in blocking buffer at 4°C, neurons were incubated with anti-rabbit Alexa-594, anti-mouse Alexa-647, or anti-sheep Alexa-488 secondary antibodies (1:500) and DAPI (0.3 µM) in blocking buffer. Coverslips were washed in PBS and mounted in Fluoromount aqueous mounting medium. Images were obtained by a Zeiss LSM 980 confocal scanning microscope equipped with a spectral detector. Images of axons and cardiomyocytes were acquired at ∼2-4 µm z-stacks at intervals of 0.25 µm using 63X objective (1.4 NA) with 3x digital zoom. Number of varicosities per 10 μm of axon length was quantified by co-localization of synaptotagmin-1/2 and TH in isolated axons contacting a cardiomyocyte using Image J. Quantification of axon growth in distal axon compartments was done by calculating integrated TH fluorescence density per unit area (ImageJ) from multiple randomly selected images from three independent experiments.

### Superior cervical ganglia injections

SCG injections were done by adapting a protocol described previously ^11^. Neonatal rats or mice (P2-P3) were anesthetized by hypothermia by laying them for 5 min on a nitrile glove placed on crushed ice. Adult animals (P30) were anesthetized by continuous inhalation of isoflurane (1-4%) for the 15 min duration of the surgery. Breathing rate of each adult animal was monitored throughout the procedure, with adjustments to anesthetic dose made as necessary. Puralube, a protective eye ointment, was applied to the eyes of adult animals. For superior cervical ganglia (SCG) injections, the area of the ipsilateral SCG was treated with depilatory cream (NAIR) for 1 min for adult animals and washed with water. The throat and neck area of all animals were swabbed with 70% ethanol and betadine before incision. An incision was made to the front of the neck, lateral to the midline using a sharp scissor. Fat, muscle, and glands were cut or moved aside to expose the SCG, residing at the juncture of the internal and external carotoid arteries. A pre-pulled microcapillary needle was used to puncture the membrane of the SCG and inject 0.3 µL of Sulfo-NHS-SS-biotin (20 mg/mL) or 0.2 µL of Rabbit anti-Flag antibody (∼0.8 mg/mL). Skin was sutured using sterile silk sutures, then swabbed with NewSkin adhesive. For adult animals, ketoprofen (25 mg/kg) was applied subcutaneously for analgesia immediately following the procedure. Animals were allowed to recover on a heating pad for 30 min. Eight hr following surgery, animals were euthanized, their SCGs, salivary glands, and irises were dissected out and subjected to streptavidin precipitation/immunoblotting or immunofluorescence.

For axotomy, after SCG injections, the SCG was separated from the carotid artery at the bifurcation to sever the post-ganglionic sympathetic nerves projecting along the carotid artery.

### Streptavidin pull-downs and immunoblotting

Tissues were lysed in RIPA buffer (1% NP-40, 0.5% sodium deoxycholate, 0.1% SDS, 50 mM TRIS, pH 7.4, cOmplete Mini protease inhibitor cocktail). After homogenization and centrifugation (13200 rpm, 4°C, 15 min), supernatants were subjected to precipitation using Streptavidin magnetic beads and immunoblotting using anti-TrkA (1:1000) or anti-NCAM (1:1000) primary antibodies, followed by incubation with anti-rabbit HRP-conjugated secondary antibody. All immunoblots were visualized using ECL Plus Detection Reagent and a ChemiDoc^TM^ Touch Imaging System equipped with a detector Cooled CCD camera (Bio-Rad).

### Immunohistochemistry in tissue sections or whole mounts

Tissue sections of SCGs (12 μm) or salivary glands (25 μm) were washed in PBS, permeabilized in PBS containing 0.1% Triton X-100, and blocked using 5% goat serum + 0.1% Triton X-100 in PBS. Sections were then incubated with mouse anti-Tuj1 (1:1000), rabbit anti-TH (1:500), or mouse anti-synaptophysin-1 (1:500) in blocking buffer overnight at 4 °C. Following PBS washes, sections were incubated with anti-rabbit Alexa Fluor-488 (1:500) anti-mouse Alexa Fluor-594 (1:500) and DAPI (0.3 µM). Sections were then washed in PBS and mounted in Fluoromount aqueous mounting medium. Images were obtained using a Zeiss LSM 980 confocal scanning microscope equipped with a spectral detector. Images were captured in z-stacks at intervals of 0.35 µm using a 63X objective (1.4 NA) with 1x digital zoom for ganglia (∼10 µm z-stack) and 3x zoom for axons (∼4-5 µm z-stack). Each channel for each image was thresholded. Number of Flag-TrkA puncta within Tuj1 positive axons was counted per 10 µm length of axons using Image J. To quantify Flag-TrkA puncta in synaptophysin-1-positive varicosities, regions of interest (ROI, 30 µm^2^) were assigned in the tissue, and the number of Flag-TrkA puncta localized within synaptophysin-1-positive varicosities in individual axons were counted using the “analyze particle” function in Image J and expressed relative to the total Flag-TrkA in the ROI.

Whole-mount immunohistochemistry to visualize nerve architecture in eye tissue, which specifically includes the iris and cornea, was done according to a protocol adapted from ^46^. Following euthanasia, animals were enucleated, and the eyes were placed in 4% PFA in PBS overnight at 4 °C. The cornea was subsequently dissected from the globe, ensuring that the iris remained attached. The part of the sclera attached to optic nerve and the lens were removed. Iris pigments were bleached by incubating the tissue in 5% hydrogen peroxide (H₂O₂) in PBS at 54°C for 4 hr. Samples were washed in PBS 3 times for 10 min each. Tissues were permeabilized by incubating with 1 % Triton X-100 in PBS at room temperature for 1 hr, then tissues were transferred to blocking buffer (5% BSA, 10% goat serum, 0.3 % Triton X-100, 0.1 % Tween-20 in PBS) for 30 min at room temperature. Samples were incubated overnight in blocking buffer with mouse anti-synaptophysin-1 (1:500), rabbit anti-TH antibody (1:500), or mouse anti-Tuj1 (1:500) at room temperature. Following PBS washes, samples were incubated with anti-rabbit Alexa-488 (1:500), anti-mouse Alexa-594 (1:500), and DAPI (0.3 µM) in blocking buffer. After washes, tissues were mounted with the iris facing upward for imaging on glass slides using Fluoromount aqueous mounting medium. Z-stack images (∼2-5 µm, 0.25 µm intervals) of nerves on iris surface were imaged using a Zeiss LSM 980 confocal microscope equipped with a spectral detector and 63x objective (1.4 NA) and 2.5x digital zoom. Image processing and analysis were performed using Image J and Imaris software. Flag-TrkA distribution in the iris was analyzed similar to that described above for tissue sections. Presynaptic varicosities were identified and quantified based on the co-localization of synaptophysin-1 and TH immunofluorescence along individual axons. Number of varicosities per 10 μm of axon length was quantified by the co-localization of synaptophysin-1 and TH in individual axons. For volumetric analysis, Imaris surface rendering tool was employed to create 3D reconstructions based on the TH fluorescence channel. Varicosity volumes were quantified by isolating TH-dense regions that overlapped with the synaptophysin-1 signal.

### Norepinephrine biosensor

Recombinant adenovirus construct expressing CMV-GRAB_NE2h_-EF1a-mCherry was generated by WZ Biosciences Inc. at a titer of 3.3×10 ^11^ pFu/mL. After co-culturing sympathetic neurons and cardiomyocytes in microfluidic chambers for 5 days *in vitro*, cardiomyocytes were infected overnight with CMV-GRAB_NE2h_-EF1a-mCherry adenoviral construct (1:1000 dilution) in culture media containing 1% FBS. After 48 hr to allow expression of biosensor, Alexa Fluor 647-conjugated Cholera Toxin Subunit B (CTB, 1 µg/mL) was added to soma + proximal axon compartments for 4–5 hr to anterogradely label axons that had projected to distal axon compartments and were contacting cardiomyocytes. Culture media was replaced with Tyrode’s solution, and culture dishes were transferred to a 37°C stage-top incubator mounted on a Zeiss LSM 980 confocal microscope equipped with a spectral detector. Imaging was done using 40X water-immersion objective lens (1.2 NA, with correction collar). Images were collected at a resolution of 512 × 512 pixels at 1 frame per 300 ms for 3 min using a fast acquisition mode to minimize photobleaching during repeated imaging. Following baseline acquisition, soma + proximal axon compartments were stimulated with nicotine (10 µM), and cardiomyocytes were re-imaged. To quantify GRAB_NE2h_ fluorescence, 4–5 regions of interest (ROIs; 9 µm²) were selected in each cardiomyocyte at axonal contact sites, identified by mCherry and Alexa Fluor 647-CTB fluorescence signals. After background subtraction, change in fluorescence intensity was calculated as: ι1F=(F-F_0_)/F_0_ where F represents fluorescence after stimulation and F₀ the baseline fluorescence. A total of 18–20 cardiomyocytes were analyzed across three independent experimental replicates.

### Cardiomyocyte beating

Cardiomyocyte beating was assessed in co-cultures as previously described ^18^. Briefly, sympathetic neurons and cardiomyocytes were co-cultured in microfluidic chambers for 7 days *in vitro*. Culture media was replaced with Tyrode’s solution, and culture dishes were transferred to a 37°C environmental chamber mounted on a Zeiss LSM 980 confocal microscope for wide-field microscopy imaging. Time-lapse images were acquired by phase-contrast microscopy using a 40X water immersion objective (1.2 NA with correction collar). Cardiomyocytes that contacted sympathetic axons were selected for imaging. Images were collected at a resolution of 512 × 512 pixels at the maximum speed allowed with a Leica DFC365 FX CCD camera at 16-bit depth and 2 × 2 binning. Time-lapse images were processed using bandpass filter in Image J (NIH) and individual cardiomyocyte beating rate was quantified. Regions of interest (ROIs, ∼3 µm length) were assigned along the plasma membrane of individual cardiomyocytes and changes in mean pixel brightness over time at individual ROIs were analyzed using Image J to quantify beat rate. At least 25 cardiomyocytes from 6 different chambers were quantified in 3 independent experiments.

### Quantification and statistical analyses

Sample sizes were similar to those reported in previous publications ^8, 20, 44, 47^. Data were collected randomly. For practical reasons, analyses of transgenic mouse lines were done in a semi-blinded manner such that the investigator was aware of the genotypes prior to the experiment but conducted the staining and data analyses without knowing the genotypes of each sample. All graphs and statistical analyses were performed using GraphPad Prism 10. Student’s t tests were performed assuming Gaussian distribution, two-tailed, unpaired, and a confidence interval of 95%. One-way or two-way ANOVA analyses with post hoc Tukey test were performed when more than two groups were compared. Statistical analyses were based on at least 3 independent experiments and described in the figure legends. All error bars represent the standard error of the mean (SEM).

## Acknowledgements

We thank Haiqing Zhao for helpful comments on the manuscript. We thank the JHU Integrated Imaging Center and JHU IBBS SOM Microscope Facility for assistance with microscopy. We thank John Kim (JHU) for use of the Gel-Doc imager. This work was supported by NIH R01 award, NS133423, to R.K., NIH R35 award, NS132153 to S.W., and a Merkin Peripheral Neuropathy and Nerve Regeneration (PNNR) Center micro grant, 24-DF/MG/301, to G.M-A.

## Author Contributions

G.M-A and R.K. designed the study. G.M.-A conducted the majority of the experiments and data analyses. M.T. assisted with confocal imaging and S.M.M, S.R, and S.W provided technical guidance and assistance with electron microscopy. G.M-A and R.K. wrote the manuscript.

## Materials and Correspondence

Correspondence and requests for materials should be addressed to R.K.

## Supplementary Materials List

A Word file with:

1. Supplementary Figures 1-6 and Figure Legends
2. Figure Legends for Supplementary Movies 1-5

## Supplementary Materials

**Supplementary Figure 1.**
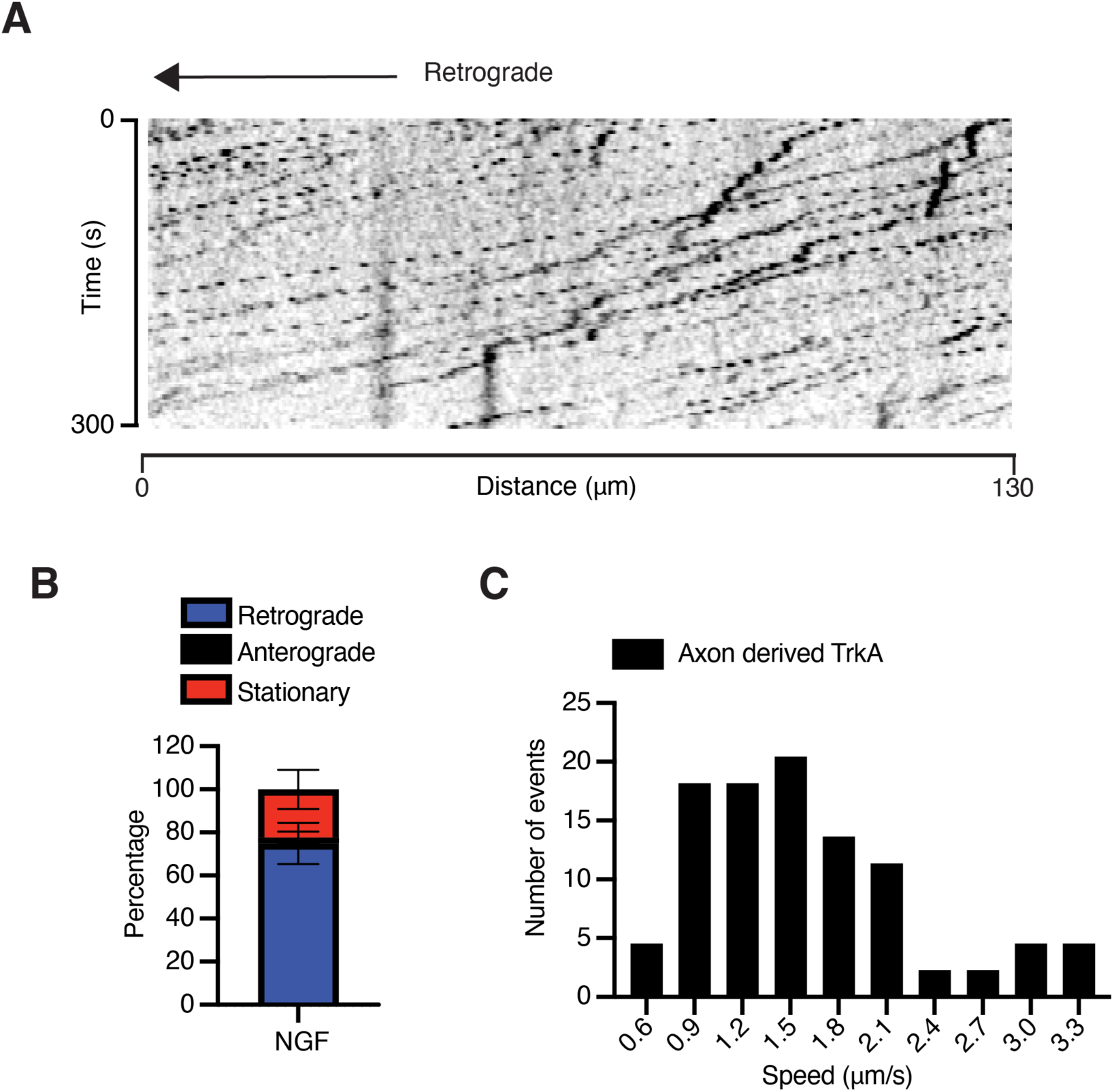
Dynamics of axon-derived TrkA receptors. Related to Figure 1. (A) Representative kymograph shows that axon-derived Flag-TrkA receptors undergo retrograde transport in a highly processive manner with few intermittent pauses. Cy3-anti-Flag antibodies were fed to distal axon compartments in microfluidic cultures of Ntrk1^Flag^ sympathetic neurons. Distal axons were stimulated with NGF (100 ng/mL, 30 min). (**B**) Quantification of directionality. (**C**) Quantification of instant speed of retrogradely transported axon-derived TrkA. Data are means ± SEM from n=3 independent experiments.

**Supplementary Figure 2.**
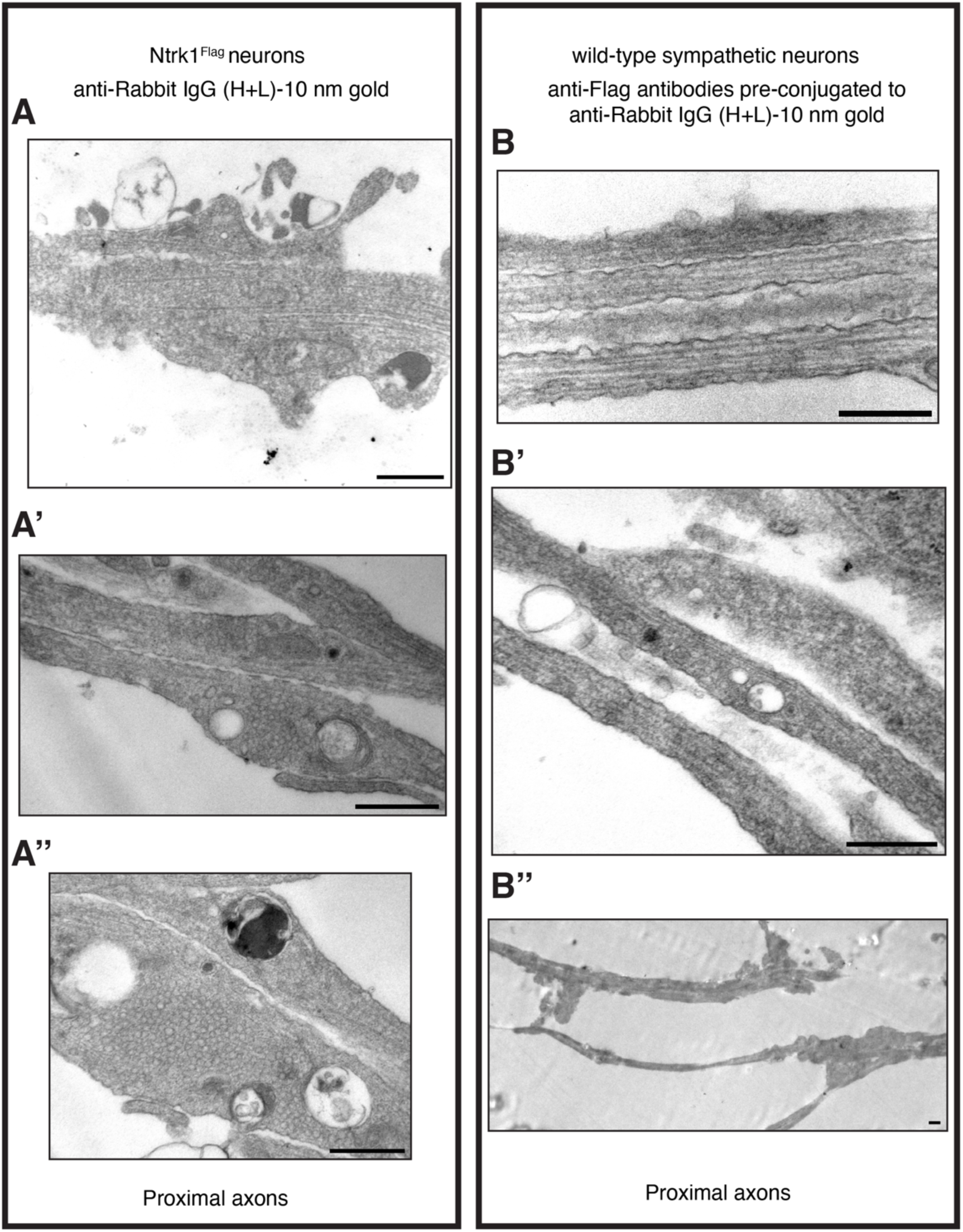
Specificity of Flag-TrkA-gold immunolabeling in EM analyses. Related to Figure 2. (**A-A”**) No electron-dense structures were observed in Ntrk1^Flag^ sympathetic neuron cultures incubated only with secondary anti-rabbit IgG-10 nm gold particles. Scale bar 200 nm. (**B-B”**) Little to no Flag-gold labeling was observed in wild-type sympathetic neurons incubated with anti-Flag antibody pre-conjugated to anti-rabbit IgG 10 nm gold particles. Representative images from proximal axon compartments are shown. Scale bar 500 nm, n=2 independent experiments for (**A,B**).

**Supplementary Figure 3.**
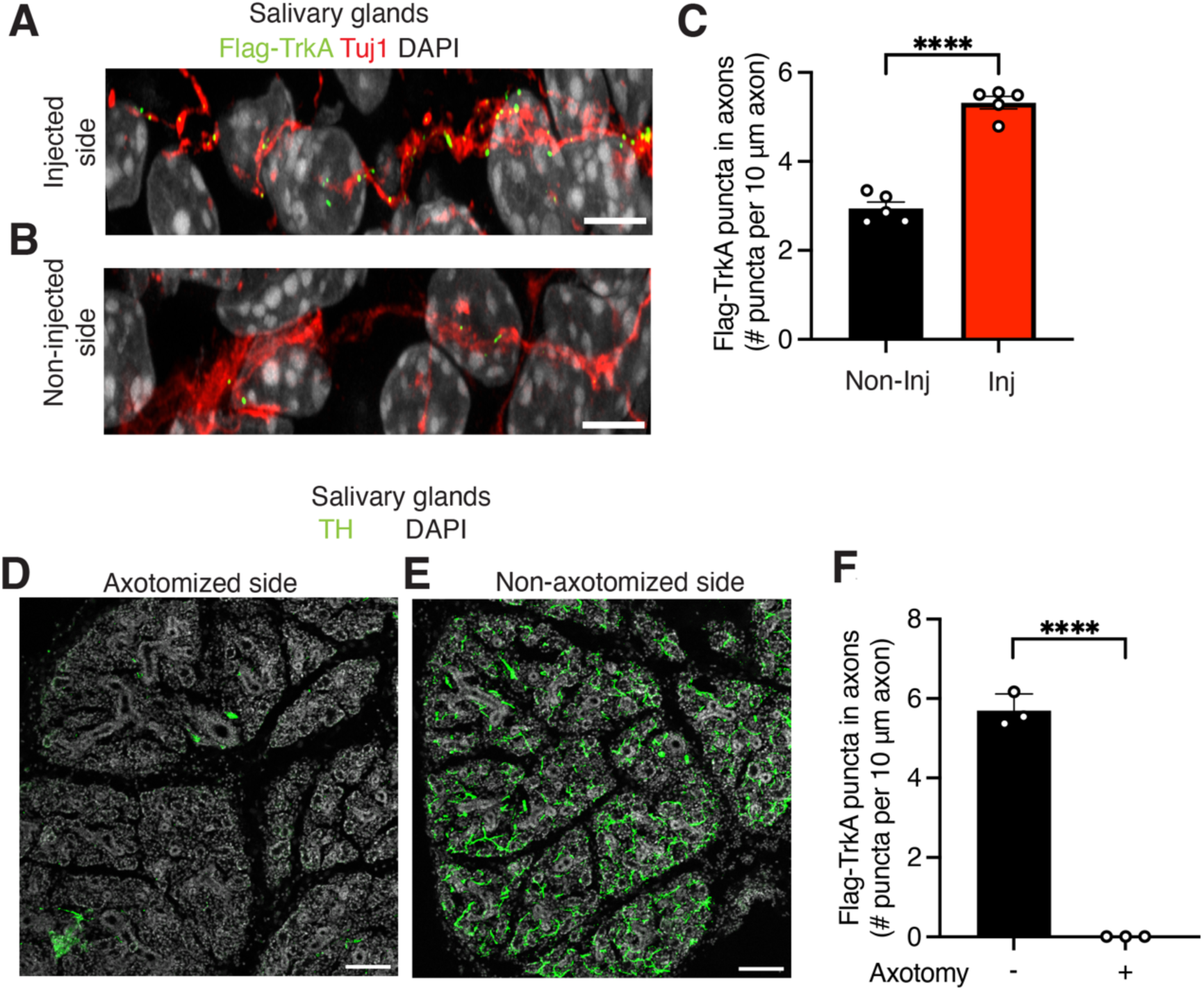
TrkA transcytosis to salivary glands is abolished by axotomy. Related to Figure 3. (**A-B**) Anterograde appearance of Flag-TrkA receptors, labeled in sympathetic ganglia, in sympathetic axons innervating salivary glands, as visualized by Flag-TrkA (green) and Tuj1 (red) immunofluorescence in P2-P3 Ntrk1^Flag^ mice. DAPI (gray). Scale bar 5 μm. (**C**) Increased Flag-TrkA puncta in salivary glands from injected side compared to non-injected side, 8 hr after SCG injection. Data are means ± SEM from n=5 animals; ****p<0.0001, t-test. (**D-E**) Unilateral axotomy of sympathetic nerves results in the loss of innervation to the salivary glands in the axotomized side, but not the contralateral side, in P2-P3 Ntrk1^Flag^ mice. Sympathetic nerves were visualized by TH immunostaining (green). Scale: 200 μm. (**F**) Anterograde appearance of soma surface labeled Flag-TrkA puncta in nerves innervating the salivary glands is abolished by axotomy. Flag and Tuj1 immunofluorescence were done 8 hr after injection of anti-FLAG antibodies into SCG in Ntrk1^Flag^ mice (P2-P3). Flag signal was compared in axotomized animals relative to the non-axotomized animals. Data: means ± SEM from n=3 animals; ****p<0.0001, t-test.

**Supplementary Figure 4.**
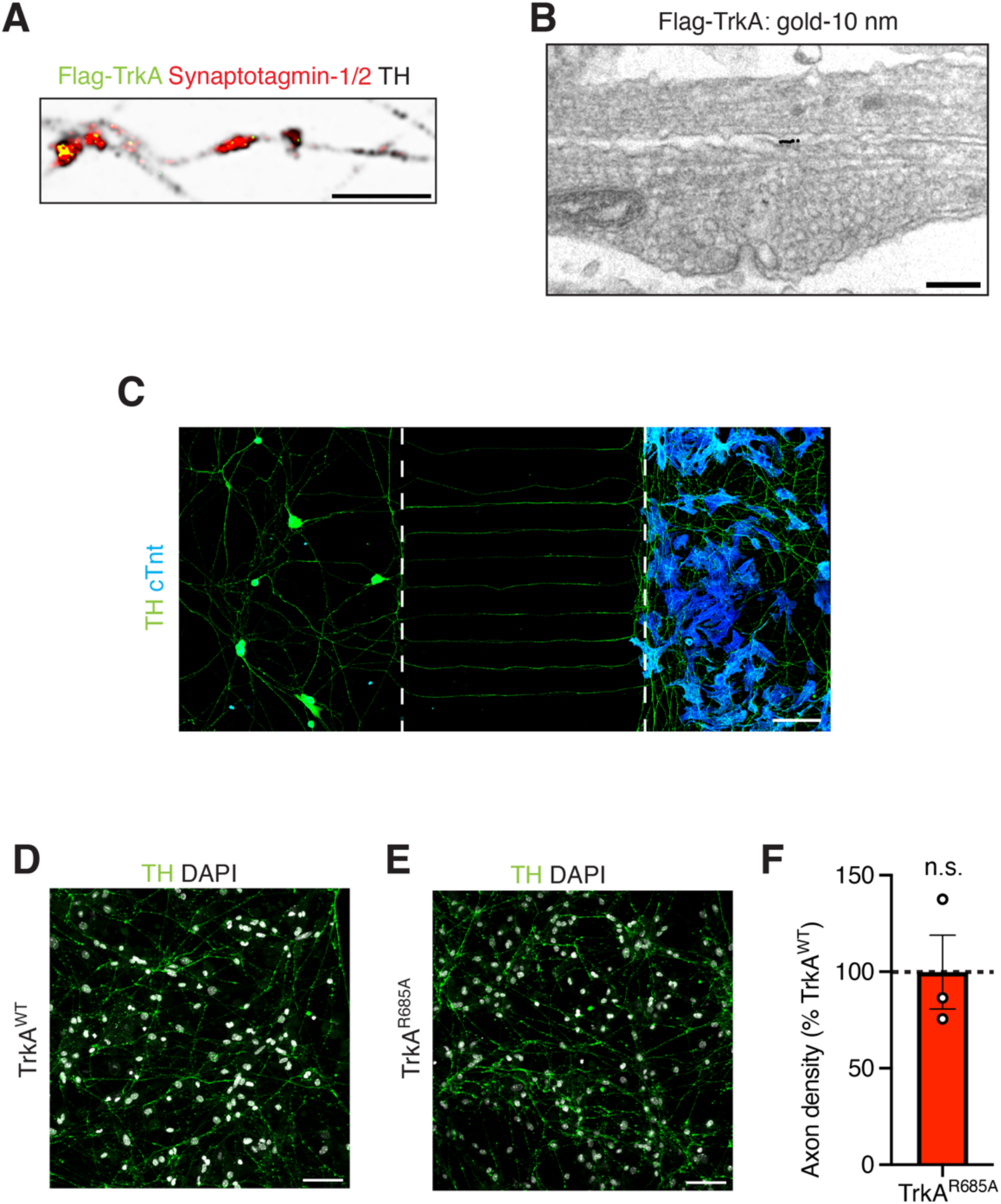
Sympathetic neuron-cardiomyocyte co-cultures to study synaptic connections. Related Toxin Figure 4. (**A**) Soma surface derived Flag-TrkA (green) accumulate in axonal varicosities marked by synaptophysin-1 (red). Flag antibodies were fed to soma compartments in microfluidic cultures of Ntrk1^Flag^ sympathetic neurons maintained by NGF (100 ng/mL). TH labeling is shown in black. Scale bar 5 μm. (**B**) EM showing Flag-TrkA gold particles at the plasma membrane in an axonal varicosity. Scale: 100 nm. (**C**) Co-cultures of sympathetic neurons and cardiomyocyte in microfluidic chambers, where cardiomyocytes were plated in distal axon compartments. Neurons are labeled by Tyrosine Hydroxylase (TH, green), and cardiomyocytes with cardiac troponin (cTnt, blue). Scale: 100 μm. (**D-E**) Axon outgrowth is similar between TrkA^R685A^ and TrkA^WT^ neurons in distal compartments in co-cultures. TH (green), DAPI (white). Scale 50 μm. (**F**) Quantification of axon density in distal axon compartments by calculating integrated TH fluorescence density per unit area (ImageJ). TrkA^R685A^ axon growth is represented as a % of the TrkA^WT^ levels Data are means ± SEM from n=3 independent experiments, n.s. not significant, t-test

**Supplementary Figure 5.**
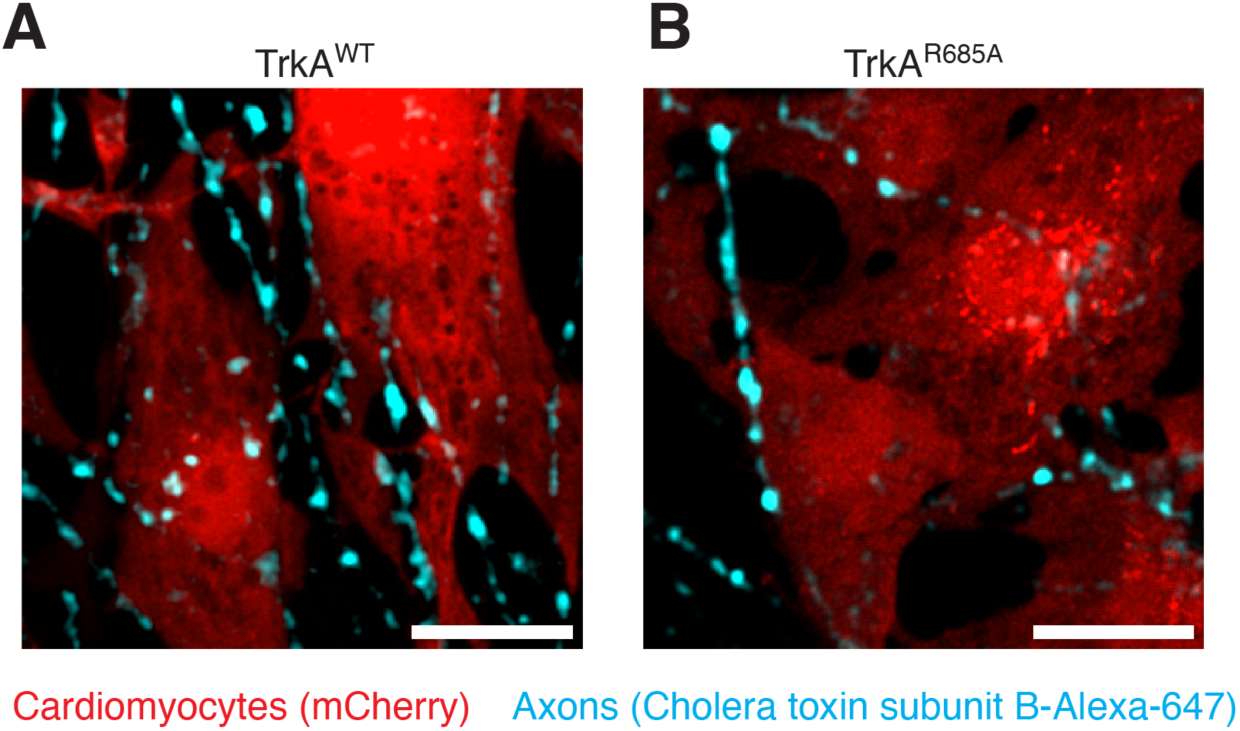
Contacts between sympathetic axons and NE biosensor-expressing cardiomyocytes. Related to Figure 5. (**A,B**) In microfluidic co-cultures, sympathetic axons make contacts with cardiomyocytes, co-expressing GRAB_NE2h_ NE biosensor and mCherry (red), in distal axon compartments. Sympathetic axons were visualized by anterograde trafficking of Alexa Fluor-647-conjugated Cholera Toxin Subunit B (CTB, cyan) added specifically to neuronal cell bodies. Scale 20 μm.

**Supplementary Figure 6.**
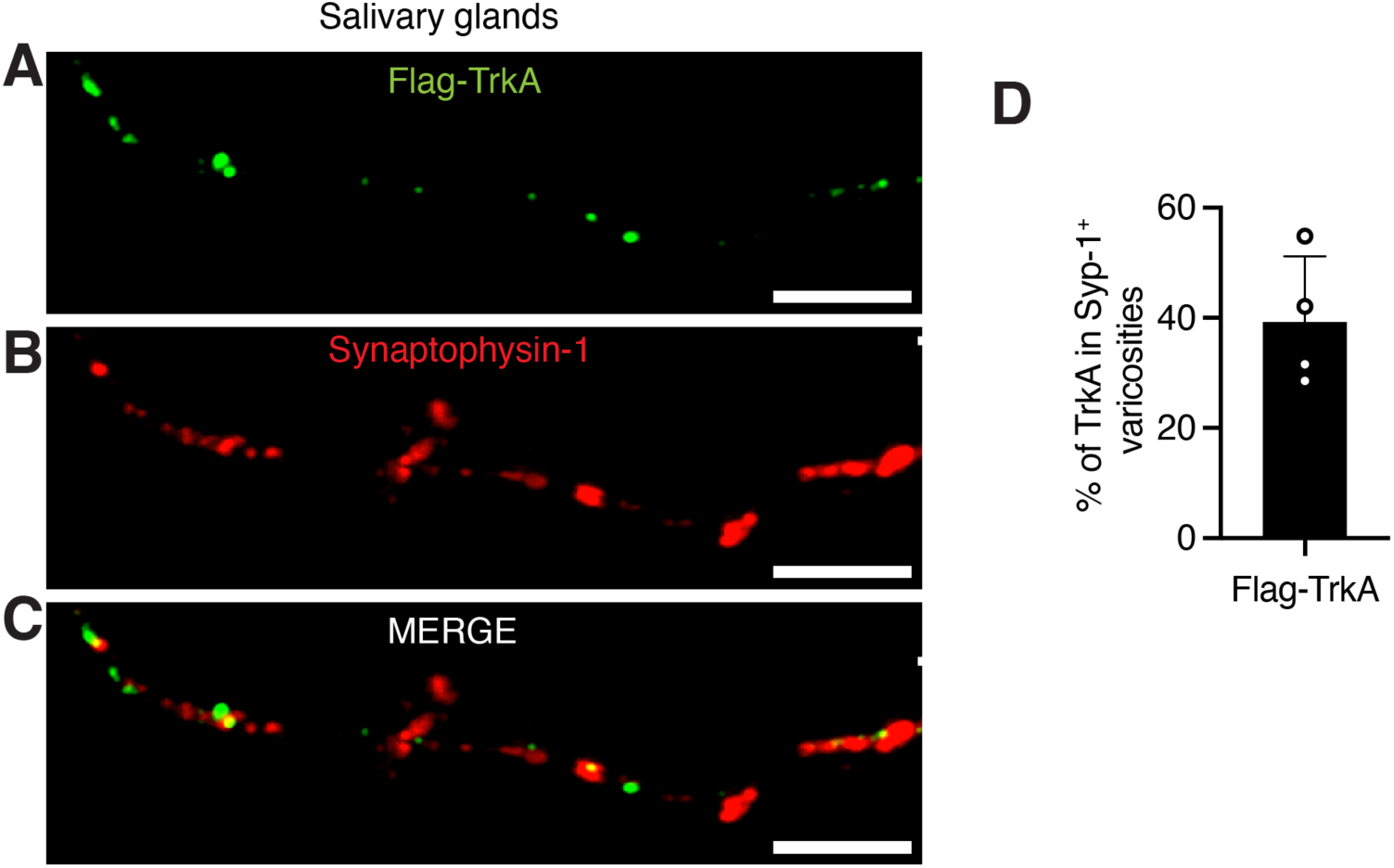
Transcytosed TrkA receptors are enriched in presynaptic varicosities in salivary glands. Related to Figure 6. (**A-C**) Transcytosed Flag-TrkA (green) appear in synaptophysin-1-positive (red) varicosities in sympathetic axons innervating the salivary glands *in vivo*. Flag and synaptophysin-1 immunofluorescence was performed in salivary glands, 8 hr after injection of Flag-antibodies into the SCG in P30 Ntrk1^Flag^ mice. Scale: 5 μm. (**D**) Flag-TrkA localized in synaptophysin-1 varicosities expressed as a percentage of total Flag-TrkA in distal axons. Data are means ± SEM from n=4 mice.

## Supplementary movie legends

**Supplementary movie 1. Live imaging of Flag-TrkA in proximal axons. Related to Figure 1**. Dynamics of Flag-TrkA transport in proximal axons after NGF (100 ng/mL) stimulation of distal axons for 30 min. Soma surface Flag-TrkA receptors were live-labeled by feeding Cy3-anti-Flag antibodies to the soma + proximal axon compartments in microfluidic cultures of Ntrk1^Flag^ neurons. Flag-TrkA particles in black. Axon is outlined in yellow. Scale bar 10 μm.

**Supplementary movie 2. Live imaging of Flag-TrkA in distal axons. Related to Figure 1**. Dynamics of Flag-TrkA transport in distal axons after NGF (100 ng/mL) stimulation of distal axons for 180 min. Soma surface Flag-TrkA receptors were live-labeled by feeding Cy3-anti-Flag antibodies to the soma + proximal axon compartments in microfluidic cultures of Ntrk1^Flag^ neurons. Flag-TrkA particles in black. Axon is outlined in yellow. Scale bar 10 μm.

**Supplementary movie 3. Live imaging of cardiomyocyte beating in monocultures. Related to Figure 5**.

Live imaging of cardiomyocytes cultured alone show spontaneous beating using phase-contrast microscopy. Scale bar 50 μm.

**Supplementary movie 4. Live imaging of cardiomyocyte beating in TrkA^WT^ sympathetic neurons co-cultures. Related to Figure 5**.

Live imaging shows that cardiomyocyte beating is enhanced in the presence of TrkA^WT^ sympathetic neurons. Co-cultures were established in microfluidic chambers, and cardiomyocytes in distal compartments were imaged using phase-contrast microscopy. Scale bar 50 μm.

**Supplementary movie 5. Live imaging of cardiomyocyte beating in TrkA^R685A^ sympathetic neurons co-cultures. Related to Figure 5**.

Cardiomyocyte beating is less pronounced when co-cultured with TrkA^R685A^ sympathetic neurons. Co-cultures were established in microfluidic chambers, and cardiomyocytes in distal compartments were imaged using phase-contrast microscopy. Scale bar 50 μm.

